# Myosin XI Interacting with a RabE GTPase Is Required for Polarized Growth

**DOI:** 10.1101/617167

**Authors:** Robert G. Orr, Fabienne Furt, Erin L. Warner, Erin M. Agar, Jennifer M. Garbarino, Sarah E. Cabral, Michelle L. Dubuke, Allison M. Butt, Mary Munson, Luis Vidali

## Abstract

The fundamental eukaryotic process of intracellular trafficking requires the interconnected activity of molecular motors trafficking vesicular cargo within a dynamic cytoskeletal network. However, in plants, few mechanistic details are known about how molecular motors associate with their secretory cargo to support the ubiquitous processes of polarized growth and cell division. A yeast two-hybrid screen of a *Physcomitrella patens* library identified a RabE GTPase as an interactor of myosin XI and subsequently demonstrated all five RabE members interact with myosin XI. Consistent with a role in polarized transport, we observed RabE at the growing cell apex and at the expanding cell plate during cell division. An in vivo cross-correlation analysis of fluorescently tagged RabE and myosin XI revealed that both species are spatiotemporally coupled, demonstrating their simultaneous involvement in polarized growth. To determine if myosin XI and RabE are directly coupled, we first computationally predicted myosin XI:RabE interface through a homology modeling-directed approach. We identified a structurally conserved residue on myosin XI, V1422, that when mutated abolished RabE binding in the yeast two-hybrid system and resulted in unpolarized plants instead of the characteristic network of filamentous cells when regenerated from single cells. Together, this work demonstrates the requirement of a direct myosin XI:RabE interaction for polarized growth in plants.

## Introduction

Polarized exocytosis requires precisely coordinated vesicle trafficking to support the essential function of polarized growth across eukaryotes, in addition to more plant-specific functions such as pathogen defense and cell wall biogenesis. (Robatzek, 2007; Bove et al., 2008; Ebine and Ueda, 2015; Bibeau et al., 2018). Plant polarized growth, or tip growth, exploits turgor pressure and active transport of secretory vesicles to drive rapid anisotropic expansion (Hepler et al., 2001; McKenna et al., 2009). At present, a mechanistic understanding of how secretory vesicles are trafficked for exocytosis in plants is lacking.

Polarized delivery of secretory vesicles is the rate limiting step for growth in plants, and is dependent upon the molecular motor myosin XI (Vidali et al., 2010; Madison and Nebenfuhr, 2013; Rounds and Bezanilla, 2013). Our traditional understanding of active transport frames myosin XI as providing the mechanical force to translocate cargo on F-actin cables, since diffusion alone is insufficient to support polarized growth (Campas and Mahadevan, 2009; Moscatelli et al., 2012; Bibeau et al., 2018). Recent work demonstrated myosin’s vesicular cargo as a site of actin assembly, suggesting myosin and vesicles function as a fluid organizing center of actin nucleation and cargo transport (Schuh, 2011; Furt et al., 2013; Pylypenko et al., 2016). Studies in both vascular and non-vascular plants demonstrated the ubiquitous requirement of myosin XI for tip growth (Madison and Nebenfuhr, 2013). Investigation of *Arabidopsis* root hairs showed that myosin XI is essential for proper movement of organelles and the organization of secretory vesicles at the cell apex (Ojangu et al., 2007; Peremyslov et al., 2008; Peremyslov et al., 2012; Park and Nebenführ, 2013). Myosin XI is necessary for polarized growth in moss (Vidali et al., 2010), and is spatiotemporally coincident with secretory vesicles at the growing apex (Furt et al., 2013). Additionally, work with chimeric myosin XIs established a strong correlation between myosin XI velocity and cell growth (Tominaga et al., 2013). Presumably, the relationship between myosin XI velocity and cell growth is a consequence of augmented delivery of myosin XI cargo; however, a direct interaction with myosin XI and its secretory cargo remained elusive.

Myosin XI is homologous to the yeast class V myosin, Myo2 in *Saccharomyces cerevisiae*. In yeast, Myo2 directly interacts with secretory vesicles using its C-terminal globular tail domain to transport vesicles to the growing bud tip and sites of cytokinesis in a stereotypical “transport cascade.” (Schott et al., 1999; Pashkova et al., 2006; Donovan and Bretscher, 2012). On the secretory vesicle, Myo2 interacts with Rab GTPases Ypt31/32 and Sec4 in a stepwise fashion to orchestrate the faithful transport of cargo to the site of exocytosis (Lipatova et al., 2008; Jin et al., 2011). Rab GTPases are dynamic molecular switches that are cyclically activated and deactivated by guanine nucleotide exchange factors (GEFs) and GTPase activating proteins (GAPs), respectively. Rab GTPases localize to specific endomembrane compartments and recruit disparate effectors when bound to GTP (Khan and Menetrey, 2013; Pfeffer, 2017). Vesicle-localized Sec4 interacts with Myo2 in a GTP-dependent manner, and both Myo2 and Sec4 interact with the plasma membrane localized hetero-octameric tethering complex, the exocyst. While the precise coordination and/or exclusivity of the interactions between myosin, Rabs, and the exocyst is unclear, together these proteins compose the terminal step of the Rab-Myosin transport cascade immediately preceding exocytosis (Lepore et al., 2018). Sec4 is homologous to the Rab8 subfamily in mammals, and despite evolutionary distance the subfamily’s interaction with class V myosin is conserved from yeast to human (Welz and Kerkhoff, 2017). An analogous Rab cascade exists in mammalian cells (Knodler et al., 2010; Mizuno-Yamasaki et al., 2012), suggesting the presence of a conserved eukaryotic polarized exocytosis mechanism.

Previous work identified RabC2a and RabD1 as effectors of *Arabidopsis* myosin XI-2 (MYA2) (Hashimoto et al., 2008). The cargo-binding domain myosin XI-2 substantially diverged from the phylogenetic grouping of XI-K and XI-C/E (Reisen and Hanson, 2007), whose sequence and subcellular localization at the growing tip more closely resembles moss myosin XI and fungal myosin V (Donovan and Bretscher, 2015; Nebenfuhr and Dixit, 2018). Furthermore, the RabC clade appears independent of the secretory pathway given RabC2a’s localization to peroxisomes (Hashimoto et al., 2008) and RabD functions upstream of the RabE clade by mediating endoplasmic reticulum to Golgi transport (Batoko et al., 2000; Zheng et al., 2005) in a manner similar to yeast Ypt1 (Vernoud et al., 2003; Wang et al., 2012). Importantly, yeast myosin V does not interact with Ypt1 (Lipatova et al., 2008). Taken together, *Arabidopsis* myosin XI-2’s characterized functions fail to adequately explain downstream polarization of secretory cargo necessary to support rapid anisotropic expansion.

The plant RabE subfamily is homologous at the amino acid level to Sec4/Rab8 (Rutherford and Moore, 2002), suggesting the presence of an analogous Rab-mediated transport model of secretory vesicles by myosin XI. Both functional and live-cell imaging studies demonstrated RabE’s involvement in anterograde transport to the plasma membrane from the Golgi, and a general requirement for plant growth and cell division (Zheng et al., 2005; Speth et al., 2009; Ahn et al., 2013). Recently, RabE was identified as functioning in concert with SCD1, SCD2, and the exocyst complex (Mayers et al., 2017). SCD1 and SCD2 form a complex with putative GEF activity for RabE, thereby suggesting an activated RabE in proximity to the exocyst. This finding tentatively places RabE in a transport pathway similar to Myo2/Sec4/exocyst (Heider and Munson, 2012); however, the presence of a molecular motor to actively transport secretory cargo remained elusive.

A novel class of plant-specific, endomembrane associated proteins were identified as myosin XI receptors, called MyoB (Peremyslov et al., 2013). The MyoB family localizes to discrete motile puncta and displays promiscuous interactions with multiple *Arabidopsis* myosin XI isoforms (Kurth et al., 2017). However, MyoB does not colocalize with secretory vesicles or any other known markers (Peremyslov et al., 2015; Kurth et al., 2017), and is absent at the cortical division site where myosin XI (Abu-Abied et al., 2018; Sun et al., 2018), Rab GTPses (Chow et al., 2008; Ahn et al., 2013), and multisubunit tethering complexes (Rybak et al., 2014) are present. Observations of robust correlations between myosin XI and vesicles in cells lacking cytoplasmic streaming (Furt et al., 2013; Bibeau et al., 2018) and well-characterized homologous myosin V-Rab interactions from yeast to human (Hammer and Sellers, 2012), strongly suggest the presence of a homologous myosin-Rab interaction in plants to mediate vesicle association and transport.

We sought to understand how Rabs coordinate with active transport to sustain polarized exocytosis in plants. The moss *Physcomitrella patens* is an attractive model to investigate this process given its lack of cytoplasmic streaming and presence of only two functionally redundant copies of myosin XI, compared to 13 isoforms in *Arabidopsis thaliana*. Use of this moss model simplifies the complexity of the confounding myosin XI background and the absence of cytoplasmic streaming reduces vesicle transport to either active or diffuse motion.

Here, we show through yeast two-hybrid, bioinformatic, and *in vivo* analyses that myosin XI specifically interacts with a RabE GTPase, and this interaction requires the myosin XI globular tail domain. RabE localizes to two regions of abundant exocytic activity, the growing cell apex and nascent cell plate. Moreover, the MyoXI:RabE interaction is required for polarized cell growth.

## Results

### Identification of RabE as a Myosin XI-Interacting Subfamily

We hypothesized that myosin XI directly interacts with secretory vesicles through a hitherto unknown receptor. As an unbiased approach to identify candidate proteins capable of binding to myosin XI’s cargo-binding domain (CBD), also known as the globular tail domain, we performed a yeast two-hybrid (Y2H) screen. In conjunction with Hybrigenics (France), we generated a high-quality *P. patens* library, which is now available for public use. To determine putative myosin XI cargo-binding partners, we screened the moss library with a bait construct composed of the canonical C-terminal myosin XI cargo-binding domain plus a small N-terminal extension into the coiled-coil domain, hereafter called MyoXI-CCT (Fig. 1a). In all studies, we used the moss myosin XIa isoform and generally refer to it as myosin XI, as our previous work showed that the two myosin XI isoforms are functionally redundant with respect to tip growth (Vidali et al., 2010). This specific MyoXI-CCT bait fragment was empirically determined through a two-step purification scheme to be required for protein solubility (Fig. S1). We screened 132 million pairwise combinations, in which positive interactions were identified by growth on synthetic defined medium (SD) lacking tryptophan, leucine, and histidine.

**Figure 1.**
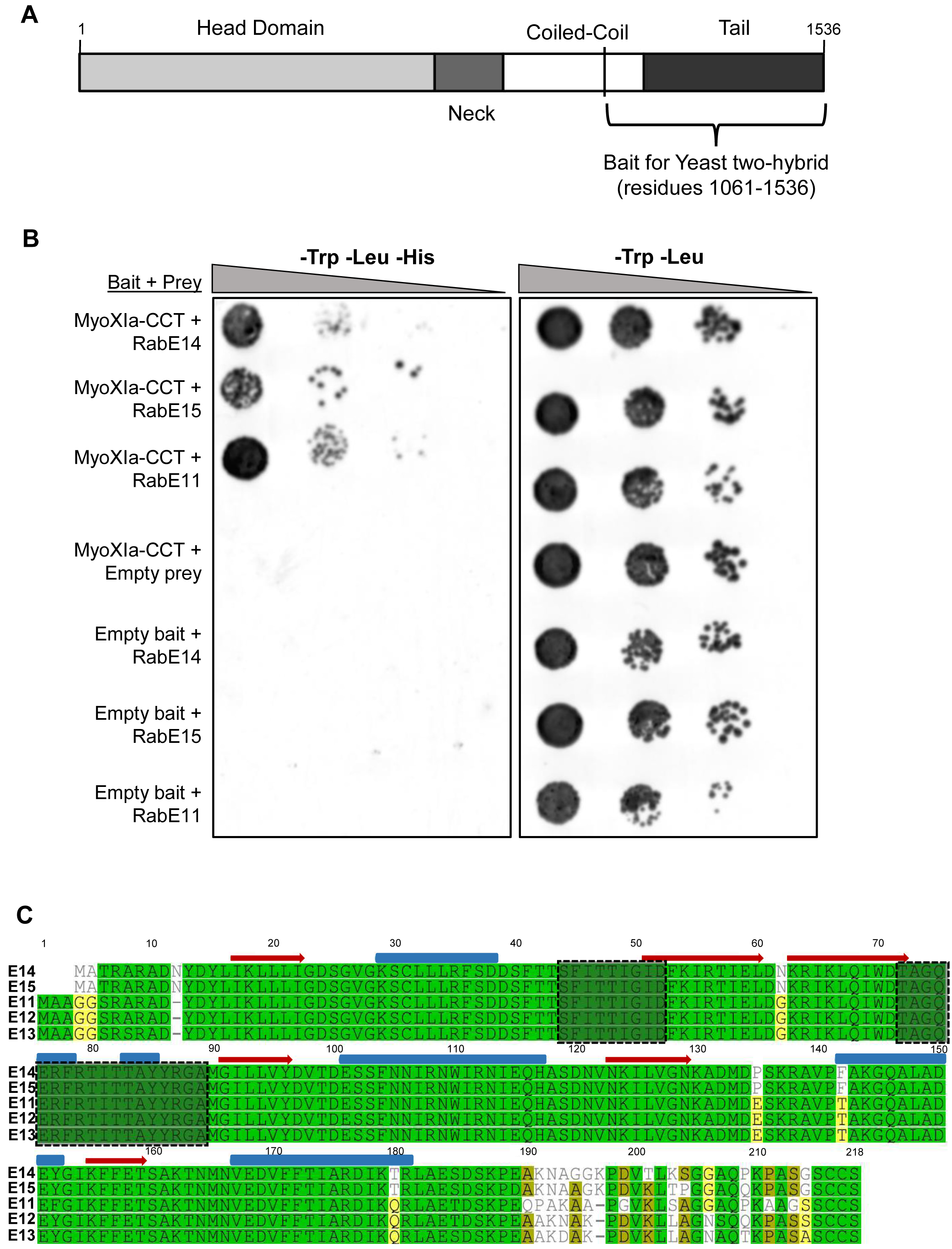
Y2H screen identified the RabE subfamily as putative myosin XI cargo-binding domain interaction partners. (A) Illustration of myosin XI’s domains and fragment (1061-1536) used to generate the Y2H bait construct. (B) Directed Y2H using the bait construct in (a) and full-length RabE14, RabE11, and RabE15 results in growth on synthetic defined (SD) medium lacking histidine. Empty bait and prey plasmids were tested with their respective partners to test for autoactivation of *HIS3*. (C) Multiple sequence alignment of the five members of the *Physcomitrella patens* RabE subfamily. Pairwise percent identity ranges from 89% (RabE14-RabE11), to 97% (RabE14-RabE15, RabE12-RabE13). Pairwise residue percent similarity is visualized by color: green=100%, yellow=60-80%, white<60%. Predicted secondary structure of active RabE14—red arrow=β-sheet, blue tube =α-helix. Switch I and II domains are outlined by the dotted line.

Our Y2H screen identified a Rab GTPase protein (Pp3c6_11710), with the interaction clone containing the full-length coding sequence of the Rab protein. As yeast two-hybrid screens can be prone to false-positives, we independently verified this putative interaction with a directed Y2H (Fig. 1b). Three independent directed Y2H experiments were performed, yielding results indistinguishable from Fig. 1. This Rab gene was initially annotated as RabE14, which we confirmed through BLAST of the *P. patens* V3.3 assembly (Lang et al., 2018). RabE14 belongs to a five-member family with high sequence conservation (89-97% identity) (Fig. 1c).

Almost all sequence diversity is located within the hypervariable C-terminal domain, while the canonical effector-determining regions localized within and near the switch regions maintain high sequence identity (Fig. 1c). Therefore, we explored the possibility that myosin XI binding is a shared function within the RabE subfamily. The most similar isoform, RabE15, and divergent isoform, RabE11, both show a positive interaction with myosin XI (Fig. 1b). We interpret this as the RabE subfamily being functionally redundant with respect to myosin XI binding. We then investigated the biological plausibility of this interaction in our moss system using the *Physcomitrella* eFP Browser (Ortiz-Ramirez et al., 2016). This expression database confirmed that all RabE paralogs are expressed in the tip-growing caulonema cell type, in which myosin XI is essential (Vidali et al., 2010).

### RabE Localizes to Sites of Polarized Secretion and colocalizes with Myosin XI

If myosin XI and RabE interact and function within the polarized transport pathway, we would expect to observe colocalization of myosin XI and RabE at subcellular locations with abundant exocytosis, such as the elongating cell apex and growing cell plate. To determine the subcellular localization of RabE we generated a constitutive expression construct of 3mCherry-RabE14 (Cherry-RabE14) and transformed this into *P. patens*. Transgenic plants that expressed the fluorescent protein and showed no growth phenotype were used for subsequent analysis. Using confocal laser scanning microscopy, we observed that Cherry-RabE14 localizes to discrete, cytosolic compartments, and predominantly accumulates at the extreme apex of growing caulonemal cells (Fig. 2a, Fig. S2b). There exists a striking resemblance between the RabE14 subcellular localization and the previously characterized apical vesicular compartment labeled by VAMP72 in *P. patens (Furt et al., 2013)*. We suspect RabE is labeling this endomembrane compartment, in addition to the free cytosolic RabE14 pool. Furthermore, this apical spot of concentrated RabE14 is dynamic, as its position and intensity vary with time (Fig. 2b, Video S1). Based on our previous data, we expected all RabE members to function redundantly. To that end, we created another RabE line using RabE12 and a different fluorophore, mEGFP. mEGFP-RabE12 displayed a similar dynamic localization to the growing cell apex (Fig. S2a), and showed no obvious growth phenotype. These data suggest that the fluorescent RabEs are functional and redundant, but this would need to be confirmed by quintuple knockouts and complementation experiments. Nonetheless, we speculated the shared apical localization could be a consequence of their myosin XI binding.

**Figure 2.**
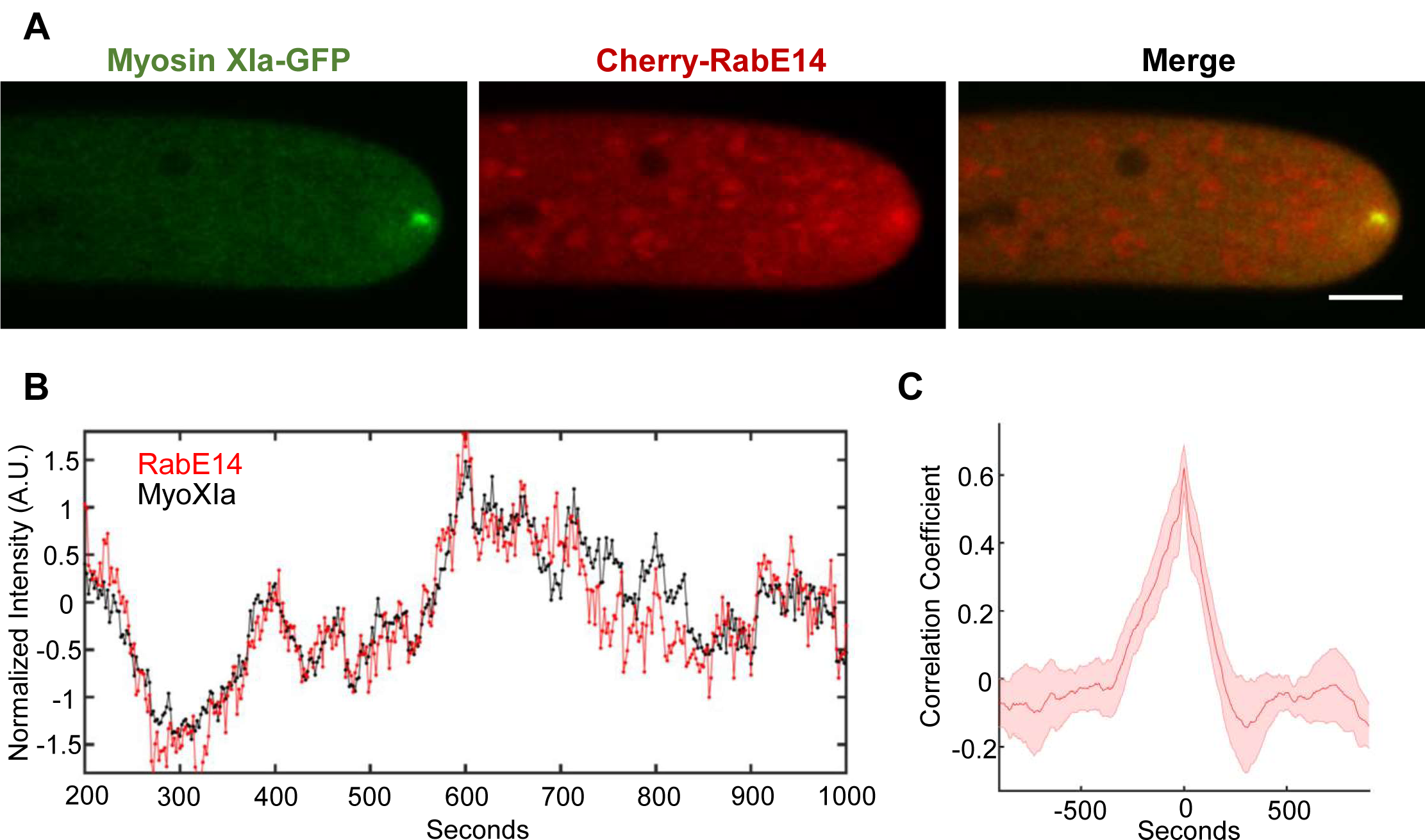
RabE localizes to growing apex and is in phase with myosin XI. (A) Subcellular localization of 3xmCherry-RabE14 and MyosinXIa-3xmEGFP by confocal microscopy. Images are maximum projections of 8 confocal slices at 1 µm spacing to visualize the apical volume. Scale bars are 5 µm. (B) Intensity fluctuations of myosin XI and RabE at the apex of the growing cell. Intensity values were obtained through kymograph analysis of time series (Video S1). (C) Maximum correlation coefficient of myosin XI and RabE intensity fluctuations. Solid line indicates the average correlation coefficient of 5 cells. The shaded region represents the standard error of the mean. Scale bar, 5 μm.

The observed apical cluster of RabE is similar to the previously observed dynamic compartment of myosin XI, F-actin, and secretory vesicles (Vidali et al., 2009; Vidali et al., 2010; Furt et al., 2013). Furthermore, myosin XI and secretory vesicles are spatiotemporally coincident, with bulk vesicle flow predominantly mediated by F-actin (Bibeau et al., 2018), presumably through myosin XI activity. This led us to hypothesize that RabE functions as a secretory vesicle receptor for myosin XI, in a manner analogous to yeast Sec4 associating with Myo2 for polarized transport (Jin et al., 2011). If true, RabE should display fluorescent fluctuations at the cell apex that are correlated with myosin XI. To test this, we generated a transgenic moss line in which 3xmCherry-RabE14 was expressed in a previously reported MyosinXIa-3xmEGFP line, where myosin Xia was tagged at the endogenous locus (Sun et al., 2018). Both myosin XI and RabE colocalized at the growing tip (Fig. 2a). Additionally, myosin XI and RabE show synchronized stochastic fluctuations of their fluorescent intensity over time within the apex of the growing moss cell (Fig. 2b). Cross-correlation analysis of the myosin XI and RabE time series reveals that these two species are in-phase (Fig. 2c). The strong observed spatial and temporal correlation suggests a meaningful biological association between myosin XI and RabE within the secretory pathway, although it cannot indicate whether it is a direct or indirect interaction.

Plants use the exocytosis machinery to transport secretory vesicles and other cargoes to build the cell plate in a manner resembling tip growth (Fendrych et al., 2010; Wu and Bezanilla, 2014; Mayers et al., 2017). Therefore, we would expect to observe myosin XI and RabE together, not only at the tip for cell growth, but also at the cell plate for cytokinesis. Consistent with recent work (Abu-Abied et al., 2018; Sun et al., 2018), we observed that myosin XI localized to the cell’s division zone immediately following nuclear envelope breakdown (Fig. 3, Video S3). RabE is noticeably absent from the division zone until approximately 9 minutes following nuclear envelope breakdown, which is consistent with the emergence of the phragmoplast (Sun et al., 2018). Upon RabE arrival at the division zone, RabE and myosin XI appear to display coordinated behavior while confined to the phragmoplast (Fig. 3, Video S3). RabE initially appears as discrete puncta and then expands outward toward the cell wall (Fig. 3, Video S3). Interestingly, unlike myosin XI, there are intense foci of RabE on the opposing plasma membranes that the phragmoplast-localized RabE grows towards. These foci appear to form via the coalescence of RabE-decorated endomembrane structures (Video S3). Upon the apparent fusion of plasma membrane-localized RabE and phragmopolast-localized RabE, coincident with myosin XI reaching the plasma membrane, the bright foci dissipate and a near homogeneous cytoplasmic signal of RabE remains (Fig. 3). Together, these observations establish RabE and myosin XI as spatiotemporally coupled in polarized transport processes. Combined with the yeast two-hybrid data, we propose that colocalization is a consequence of a direct interaction between myosin XI and RabE.

**Figure 3.**
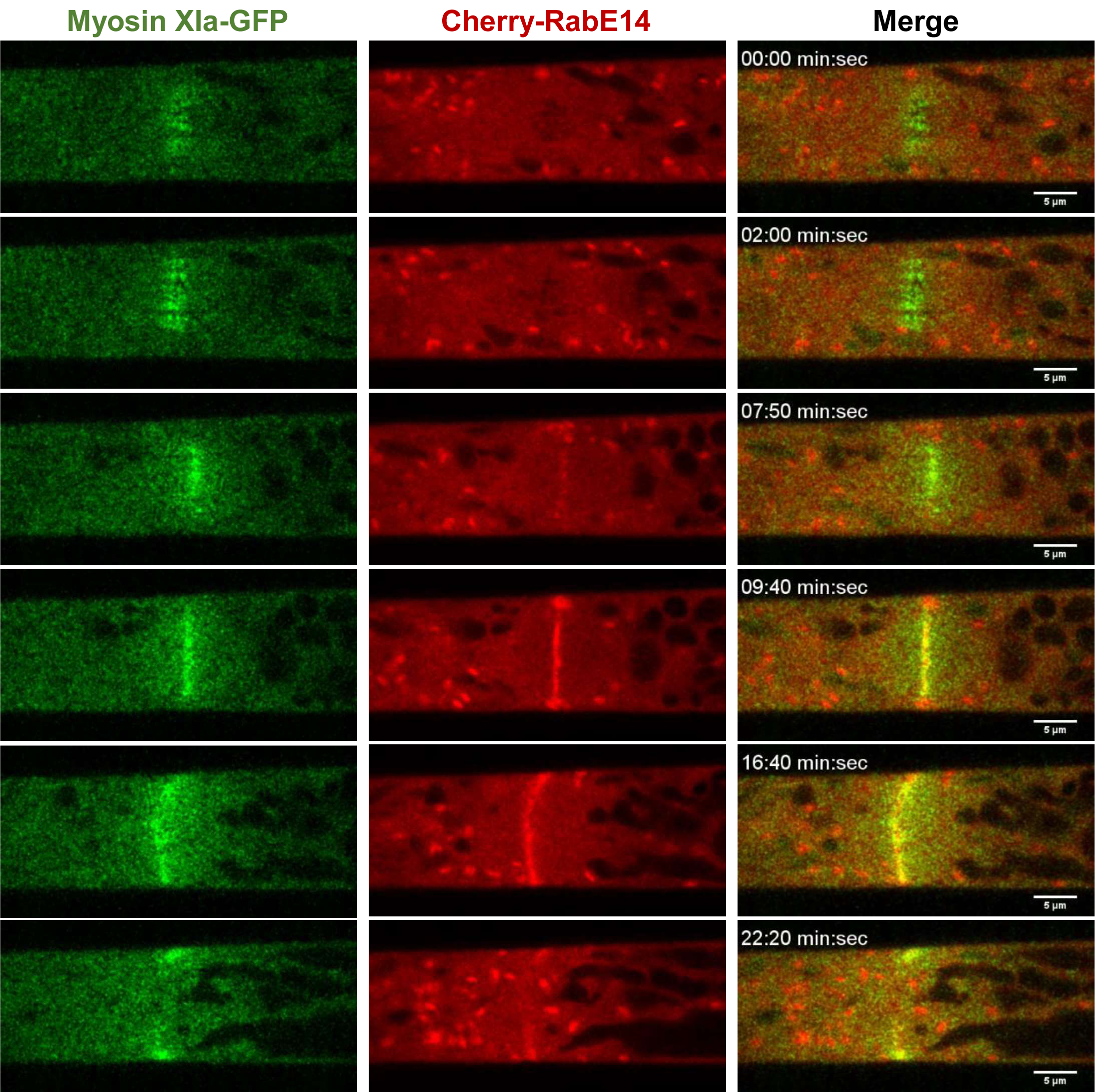
RabE co-localizes with myosin XI at the growing cell plate. An apical caulonemal cell of the double myosinXI-GFP and Cherry-RabE14 line was imaged during cell division approximately 1-minute post nuclear envelope breakdown. Scale bar, 5 µm.

### RabE Phylogeny and MyoXI:RabE Interface Prediction

To gain molecular insights of the MyoXI:RabE interaction, we hypothesized that the binding interface is structurally homologous that of to the well characterized class V myosin binding to Rab8/Sec4 (Goldenring et al., 2012; Welz and Kerkhoff, 2017). We reasoned that the stereotypical Rab effector-binding regions will be conserved across eukaryotes. Rabs undergo a dynamic GTP-dependent rearrangement of two structural regions, named “switch-I” and “switch-II,” which defines their “active” GTP-bound state and effector recognition (Stroupe and Brunger, 2000; Khan and Menetrey, 2013). To investigate the conservation within the Rab effector-binding region, we performed a multiple sequence alignment of moss RabE14, Sec4 from *Saccharomyces cerevisiae*, and Rab8a from *Homo sapiens* (Fig. 4a). RabE14 is 52% identical to Sec4 and 62% identical to Rab8a throughout the entire sequence. As expected, the most conserved residues are grouped in the switch-I, interswitch, and switch-II regions, with the greatest sequence divergence observed within the hypervariable C-terminal domain (Fig. 4a). Given the extent of conservation, we speculated that structural models of the myosin XI-tail and RabE14 that are based on the well-characterized Sec4 and Myo2 interaction could assist in mapping the functional interface.

**Figure 4.**
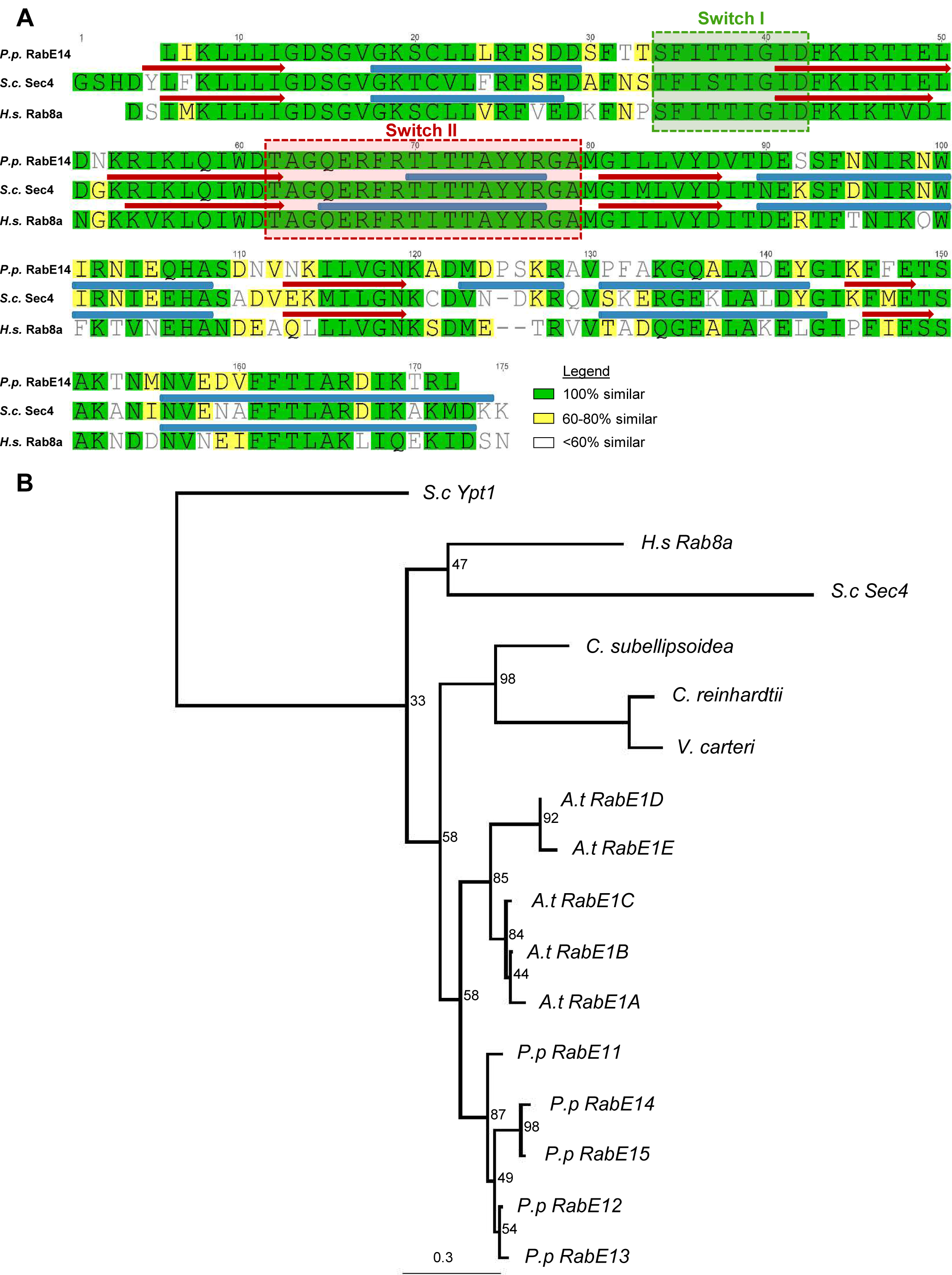
Multiple sequence alignment and phylogenetic tree of RabE14 homologs. (A) Multiple sequence alignment of *Physcomitrella patens (P.p.)* RabE14, *Saccharomyces cerevisiae (S.c.)* Sec4, and *Homo sapiens (H.s.)* Rab8a. Scondary structure (red arrow=β-sheet,blue tube=α-helix) of Sec4 and Rab8a was manually annotated based on their crystal structures, PDB IDs 1G17 and 4LHW, respectively. (B) ML phylogenetic tree of RabE subfamily members from select *Viridiplantae* species (*Coccomyxa subellipsoidea, Chlamydomonas reinhardtii, Volvox carteri, Arabidopsis thaliana, Physcomitrella patens*) and homologous Rab8 members from yeast and human. *Saccharomyces cerevisiae* Ypt1 (Rab1) was used as the outgroup. Numbers indicate support values.

We generated a phylogenetic tree to gain insight into the extent of evolutionary conservation between Rabs within same putative subfamily, and used a homology-directed approach to construct a putative MyoXI:RabE14 interaction surface. We selected representative *Viridiplantae* species and their annotated RabE subfamily members from Phytozome (phytozome.jgi.doe.gov), along with well-characterized Rabs from yeast and human and generated a maximum likelihood (ML) phylogenetic tree (Fig. 4b). This tree shows a partitioning of the plant RabE subfamily with the human Rab8a and yeast Sec4, as well as the moss and *Arabidopsis thaliana* RabEs forming discrete monophyletic groups with a shared origin (Fig. 4b).

To identify the putative interface between RabE14 and myosin XI, we first generated homology models of *P. patens* myosin XI CBD and RabE14 using yeast Myo2-CBD and Sec4-GTP as templates (Fig. 5a). Our homology models of MyoXI-CBD and RabE14 displayed good stereochemical quality, as evaluated by Ramachandran plot (Fig. S3), and average root-mean-squared deviations of 0.291 Å and 0.134 Å, respectively (Fig. S3b, d).

**Figure 5.**
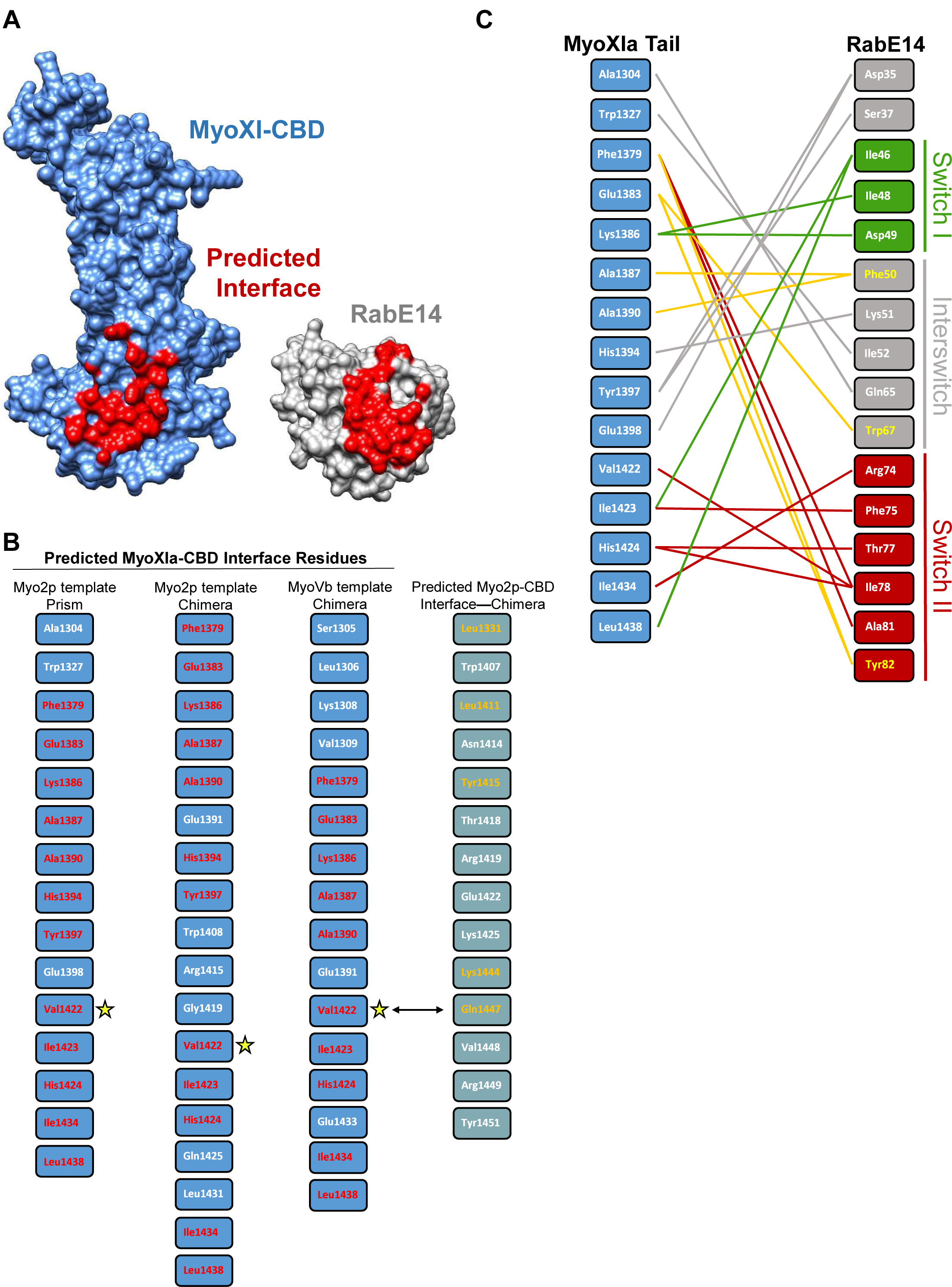
Prediction of Myosin XI-CBD and GTP-bound RabE14 binding interface. (A) Predicted MyoXI:RabE interface using homology modeling and structural superposition. (B) Predicted interface residues for MyoXI-CBD using different templates and methods results in similar interface profiles. V1422 (star) was shared between all interface predictions and aligns with known interaction residue Q1447 from S.c. Myo2p. All experimentally identified yeast Myo2p interaction residues were captured with structural superposition as described in methods. (C) Contact map of modeled myosin XI:RabE14 (−17.8 kcal mol-1) determined by template-based docking algorithm PRISM (Tuncbag et al., 2011). Hydrophobic triad residues and their putative contacts colored in yellow. Colors of lines representing contacts reflect the domain within RabE14 that the contact belongs. Green=Switch I, Red= Switch II, Grey=Interswitch, Yellow=Hydrophobic triad.

We leveraged these homology models to computationally predict the binding interface between myosin XI and RabE14. To increase the probability of identifying biologically relevant residues at the interface, we employed two template-based approaches. In the first approach, we structurally superimposed our homology models with a co-crystal structure of myosin V binding to an active, GTP-bound Rab (PDB ID: 4lx0). We considered residues containing atoms within an intermolecular distance of up to 1 Å as the putative MyoXI:RabE interface (Fig. 5a). To validate the method, we used yeast Myo2p and successfully recapitulated known Sec4 interaction residues (Fig. 5b). We also employed the template-based docking algorithm, PRISM (Tuncbag et al., 2011; Baspinar et al., 2014) to reduce the possibility of overfitting to a rigid co-crystal structure as this algorithm incorporates flexible refinement to resolve steric clashes.

We visualized the predicted residue interactions with a contact map (Fig. 5c). At the amino acid level, almost all predicted associations occur within the switch-I, interswitch, switch-II region (Fig. 5c). The hydrophobic triad, a major structural determinant of effector specificity (Merithew et al., 2001), is present in this predicted interface network. Importantly, both interface prediction methods and myosin templates identified the myosin XI residue V1422 as a putative interface residue (Fig. 5b). V1422 of myosin XI aligns directly with Q1447 of Myo2p (Fig. 5b); Q1447 was previously determined to reside in the yeast secretory vesicle binding site (Pashkova et al., 2006) and later shown to directly associate with the Rabs Ypt31/32 (Lipatova et al., 2008) and Sec4 (Jin et al., 2011). Altogether, these results support a remarkable degree of structural conservation, and suggest that RabE14 functions as a myosin XI receptor on secretory vesicles.

### Structure-based Mutants Disrupts the MyoXI:RabE Interaction

Prediction of the putative MyoXI:RabE interaction identified several candidate residues on myosin XI-CBD that could mediate an interaction with RabE. Residue V1422 was an enticing candidate given its structural alignment with Q1447 in the yeast Myo2p-CBD (Fig. S4). Residue Q1447 was previously shown to be necessary for Rab GTPase binding, specifically Ypt31/32 and Sec4 (Pashkova et al., 2006; Lipatova et al., 2008; Jin et al., 2011). To test if V1422 is required for myosin XI-CBD to interact with RabE14, we performed a directed Y2H assay in which V1422 was mutated to arginine. We generated an additional myosin XI-CBD mutant, V1418R, to probe this local region. Consistent with our interface prediction, both mutants eliminated the interaction with RabE14 (Fig. 6a). However, it appeared that V1418R imparted a slightly toxic effect, as observed by reduced growth on the control SD plate (Fig. 6a). Furthermore, the myosin XI-CBD homology model predicts that the V1418 side chain is buried, relative to V1422 (7.7 Å^2^ and 36.2 Å^2^ solvent accessible surface area, respectively). The buried attribute of V1418 suggests that the observed loss of interaction in the V1418R mutant may be a consequence of misfolding or instability of the protein, whereas the V1422R mutant specifically destabilizes the interface with RabE14.

**Figure 6.**
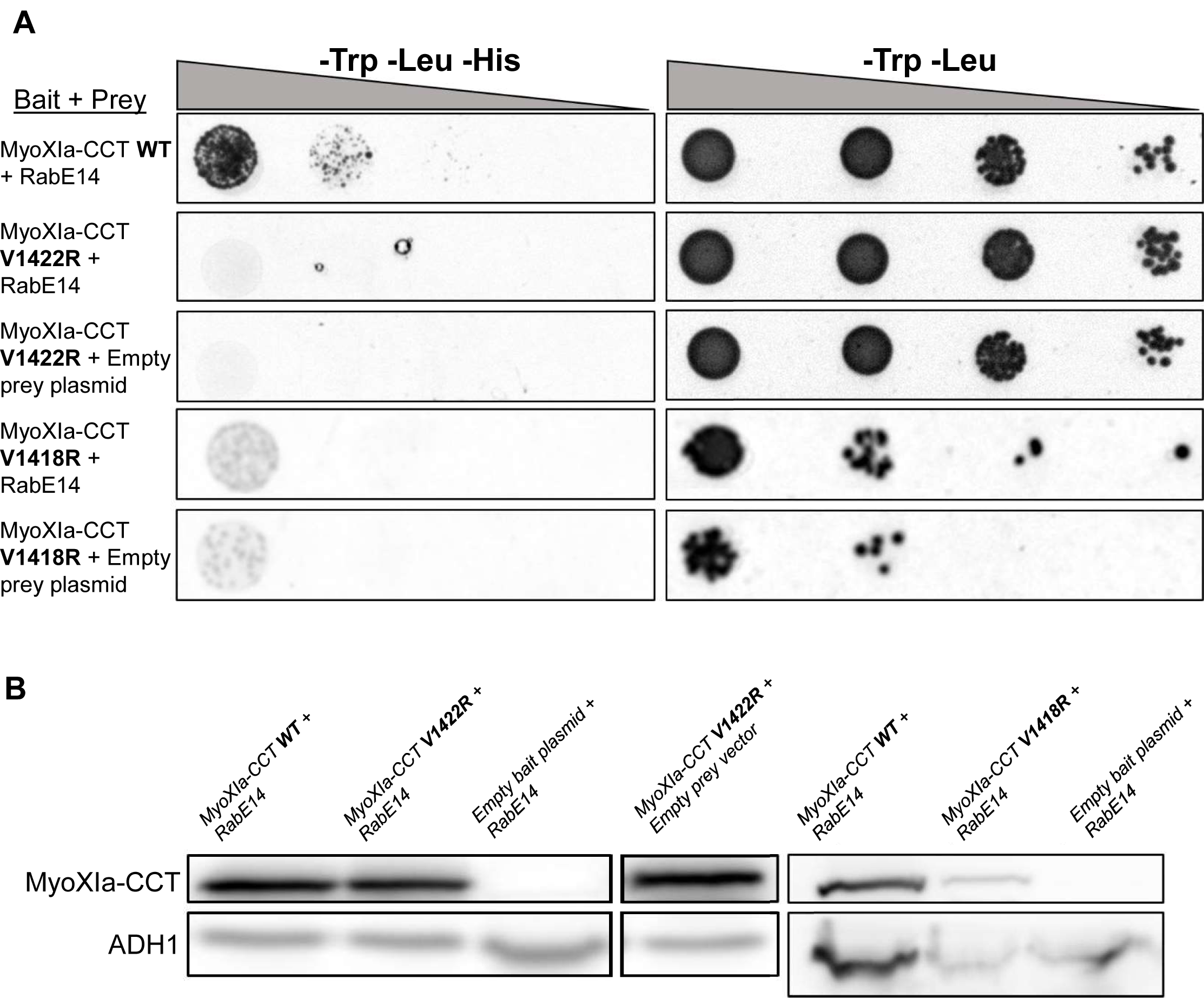
A single amino acid in Myosin XI cargo-binding domain abolished interaction with RabE14. (A) Directed Y2H with the WT bait fragment from (Fig. 1) and bait fragments containing mutations in the myosin XI cargo-binding domain that are required for polarized growth and predicted to interact with RabE14. All yeast strains were grown on the same SD plates, but rearranged for clarity. (B) Strains from (a) were grown in SD medium lacking trp and leu to an OD600 of 1 and assayed for expression of myoXIa-CCT via western blot. Total extracted yeast protein was blotted using a myoXI-CCT antibody and ADH1 as a loading control. The condition MyoXIa-CCT V1422R+empty bait plasmid belongs to the western blot shown on the left but was rearranged for clarity.

One possibility that would explain the absence of an interaction of the myosin XI-CCT mutants would be insufficient expression of the myosin XI-CCT proteins; therefore, we tested the expression in each Y2H yeast strain. Myosin XI-CCT expression was assayed using an antibody raised against the moss myosin XI-CCT (Fig. S1). The empty bait plasmid shows no myosin XI-CCT expression, whereas all other groups display comparable myosin XI-CCT expression when normalized to the loading control (Fig. 6b). The presence of V1422R at a comparable protein level to wild-type myosin XI-CCT, and the absence of a RabE14 interaction support our hypothesis that the V1422R mutation specifically abolishes RabE14 binding. Altogether, these results indicate that RabE14 directly binds to myosin XI via its cargo-binding domain, and that the V1422 residue is essential for the MyoXI:RabE interaction.

### Disruption of MyoXI:RabE Interface in vivo Impairs Polarized Growth

The MyoXI:RabE interaction is predicted to be a fundamental component of the polarized transport process. Therefore, disruption of this interaction should manifest as polarized growth defects and/or loss of polarized growth. To disrupt the MyoXI:RabE interaction, we mutated several residues, including V1422. These residues were chosen based on the modeled interface and structural agreement with known yeast Myo2 residues that interact with Sec4 (Fig. 5, Fig. S4)(Jin et al., 2011). The myosin XI-CBD mutants were cloned into the full-length myosin XI coding sequence (myosin XI containing the motor/neck/coiled-coil and cargo-binding domain) with only a single point mutation each. To test the capability of these myosin XI-CBD mutants to function in polarized growth, we used a rapid RNAi-based polarized growth assay that allows robust screening of hundreds of actively silenced moss plants (Vidali et al., 2007). Previous work demonstrated that silencing endogenous myosin XI at a 5’ UTR regions results in a loss of polarized growth; this loss is rescued upon exogenous expression of the wild-type myosin XI coding sequence (Vidali et al., 2010). Therefore, we tested if expression of any of the mutant myosin XI coding sequences could functionally complement the polarized growth phenotype, by co-transforming the myosin XI silencing construct along with a myosin XI mutant into moss protoplasts, followed by 1-week regeneration and visualization.

To characterize the polarized growth phenotype of the mutant myosin XI moss, we quantified two morphological parameters of the plant, area and solidity (Vidali et al., 2010). Solidity is the ratio of the plant area to the convex hull area, which is used as a morphological metric of plant polarization. We extracted these parameters by imaging actively silencing plants one-week post-transformation (Fig. 7b). As expected, silencing of endogenous myosin XI results in small and round plants, as measured by significantly reduced area and solidity, respectively (Fig. 7c, d). All the mutant plants show a statistically significant increase in solidity compared to control RNAi, however, the Y1384, V1422R, and V1418R mutants show a statistically significant decrease in normalized plant area with respect to the silencing negative control and wild-type rescued plants (Fig. 7c,d). Nevertheless, the F1379R mutant plants display similar polarization to the wild-type rescue (Fig. 7b) and share a small and no significant difference average plant area (Fig. 7d). The V1422R and V1418R mutants are morphologically equivalent to silencing myosin XI alone, and Y1384 shows an intermediate phenotype. The dramatic loss of a polarized growth phenotype observed in myosin XI-CBD mutants demonstrates that these individual residues are required for proper functioning of MyoXI in polarized growth.

**Figure 7.**
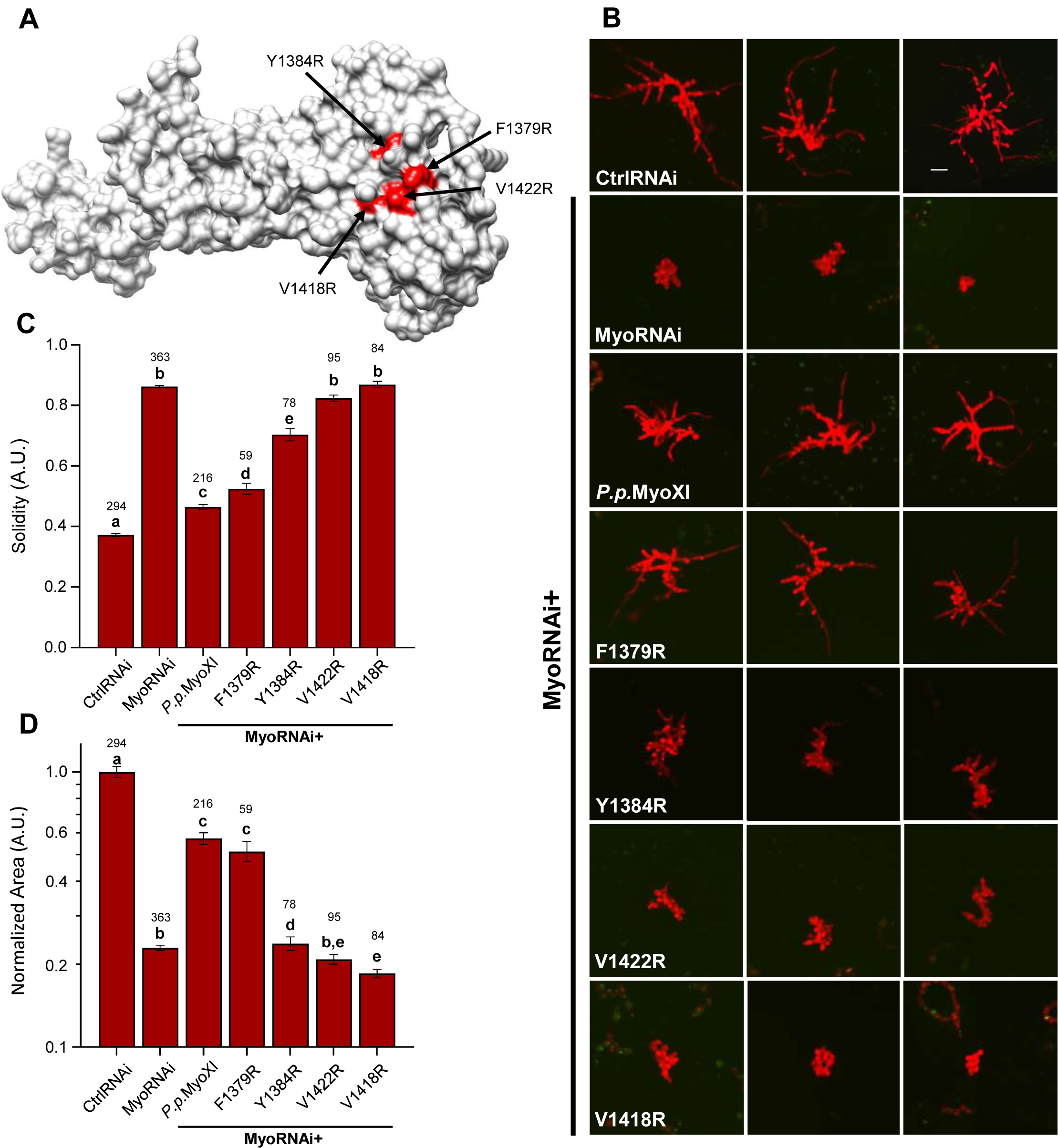
Structure-guided mutagenesis of myosin XI-tail revealed polarized growth mutants. (A) Homology model of P.p. myosin XI-tail is shown, with the residues predicted to mediate RabE interactions in red. (B) Representative 1-week old RNAi plants—all plants except ‘CtrlRNAi’ are silencing expression of both endogenous myosin XIs. All plants except ‘MyoRNAi’ were co-transformed with the myosin XI silencing construct and another construct expressing either WT myosin XI or mutant myosin XI. Bar = 100 µm. (C-D) Quantification of the morphological parameters of solidity and area extracted from images in (b). Error bars indicate standard error of the mean. Shared letters above the bars show those experimental groups that cannot be statistically distinguished. Statistical significance was determined by a one-way ANOVA-Tukey (P<0.01). Numbers above the letters indicate number of plants analyzed per condition.

We wanted to distinguish whether our observed mutant phenotypes were a consequence of the introduced mutation on the interaction with RabE, or if the mutations led to decreased protein expression and/or stability. We tested this by creating fluorescent GFP fusions of the myosin XI proteins, with GFP as a reporter for protein expression. We transformed these GFP-myosin XI constructs into wild-type protoplasts and imaged them after one day of regeneration (Fig. 8a). This approach revealed that under these conditions, the myosin XI mutants expressed at levels comparable to the wild-type myosin XI (Fig. 8b).

**Figure 8.**
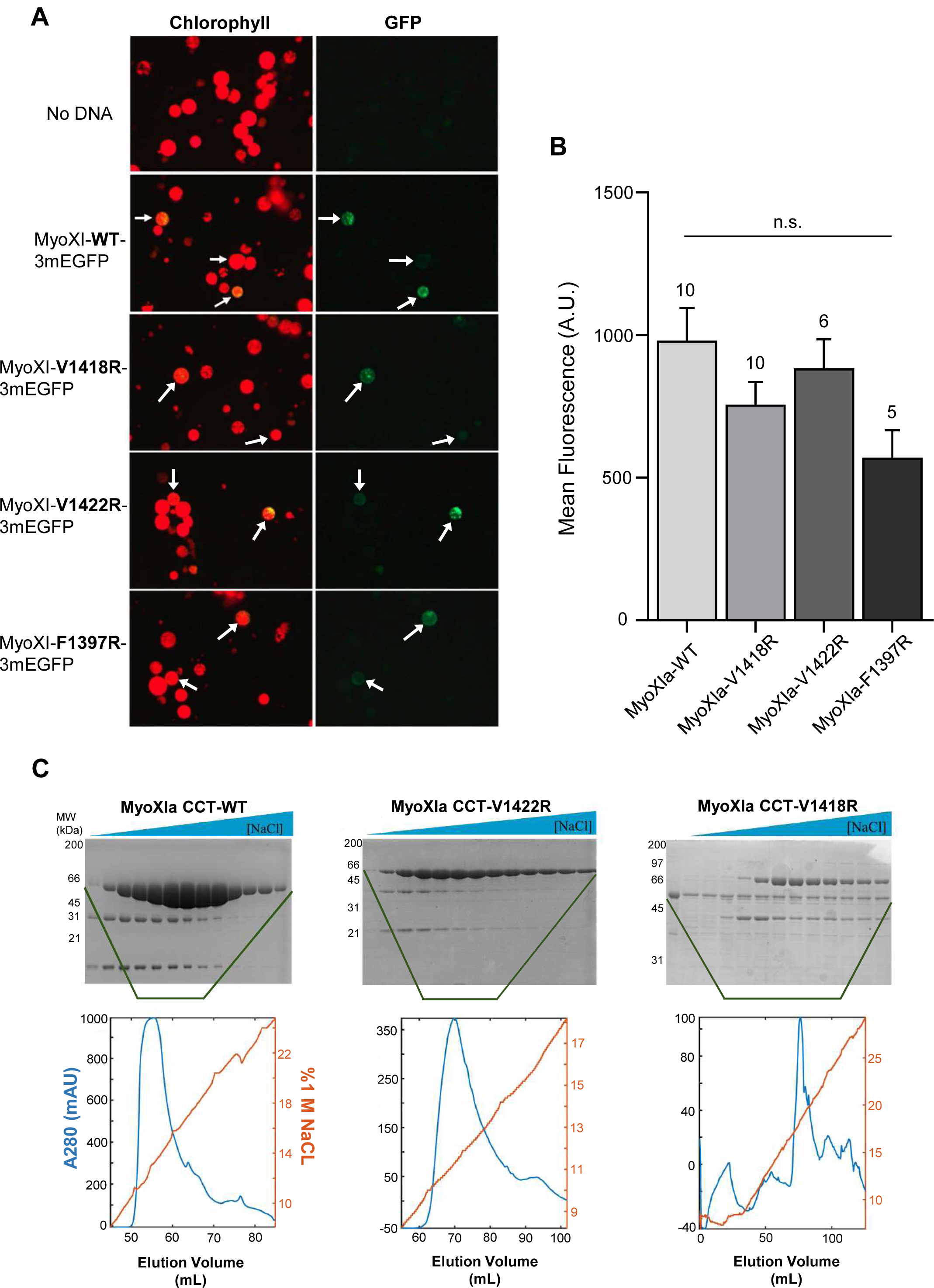
Expression of Myosin XI tail mutant proteins showed varying levels of solubility and/or stability. (A) Representative fluorescent images of wild-type moss protoplasts one day post-transformation with the indicated construct. (B) Quantification of expression for each fluorescent myosin XI fusion construct for images obtained in (a). Within a given condition, the mean fluorescence across all GFP positive protoplasts for a single independent transformation was treated as one sample. We plotted the mean of these independent sample means and failed to distinguish a statistically relevant difference, as tested by a one-way ANOVA (P<0.05). Error bars indicate the standard error of the mean and numbers above the SEM indicate the number of independent experiments. (C) SDS-PAGE gels of the elution fractions for WT and mutant myosin XI proteins following anion exchange chromatography. Each lane of the SDS-PAGE corresponds to fractions eluted from the column, with the absorbance at 280nm and percentage of 1 M NaCl plotted below.

As we had observed an apparent cytotoxic effect of the V1418R mutant in Y2H (Fig. 6a), we speculated this toxicity was due to misfolding of this mutant, perhaps a consequence of destabilizing the predicted buried side chain of V1418. To biochemically test the stability of the mutants V1422R and V1418R, we recombinantly expressed them in *E. coli*, and purified them using a two-step affinity and anion-exchange protocol. The anion-exchange chromatograms and SDS-PAGE of the purified myosin XIs show the stability of wild-type and V1422R myosin XI, but lower expression and solubility of V1418R under identical conditions (Fig. 8c). Together, these results indicate that defects due to mutation of V1418 are likely due to destabilization of myosin XI, whereas V1422 is required to mediate the polarized growth process, and the V1422R mutation specifically abrogates its biological function through defective binding to RabE proteins.

## Discussion

Our studies identified a critical interaction between the molecular motor myosin XI and the RabE GTPase subfamily. Despite multiple independent lines of research demonstrating the requirement of myosin XI and RabE for polarized trafficking, and apparent homology to myosin V and Sec4, (Zheng et al., 2005; Peremyslov et al., 2008; Speth et al., 2009; Vidali et al., 2010; Peremyslov et al., 2012; Park and Nebenfuhr, 2013) direct evidence supporting a myosin:Rab-driven transport model, as in budding yeast, was lacking (Peremyslov et al., 2013; Ryan and Nebenfuhr, 2018). The presence of a conserved eukaryotic myosin:Rab interaction in plants was previously challenging to identify. We attribute this difficulty to the expansion of the myosin XI family in vascular plants, background cytoplasmic streaming that confounds observations of vesicle transport, and existence of a promiscuous, plant-specific interaction between myosin XI and a unique endomembrane compartment (Kurth et al., 2017). By exploiting the reduced gene family of myosin XI and absence of cytoplasmic streaming in the moss *Physcomitrella patens*, in combination with Y2H and homology modeling, we identified functional homology between myosin XI/myosin V and RabE/Sec4.

Our Y2H screen used the myosin XI cargo-binding domain from the moss *P. patens* as bait, and isolated one member of the RabE subfamily, RabE14, and subsequent Y2H with other RabE members led us to conclude that binding of myosin XI is a general property of the RabE subfamily (Fig. 1b). Furthermore, generation of stable moss lines containing different RabE paralogs fused to either GFP of mCherry displayed similar dynamic subcellular localization (Fig. 2, Fig. S2). Our conclusion of the functional redundancy of the RabE subfamily is consistent not only with initial predictions (Pereira-Leal and Seabra, 2001; Rutherford and Moore, 2002), but subsequent experimental evidence demonstrating conserved RabE effector binding (Camacho et al., 2009; Mayers et al., 2017).

While observing RabE during cell division, we noticed that in addition to RabE co-localizing with myosin XI on the growing cell plate there existed RabE-labeled intracellular organelle-like structures. Given the high degree of sequence conservation between *P. patens* and *A. thaliana* RabE (∼80% identical), especially within the canonical effector binding regions, we infer that discoveries in both systems are translatable. Using that assumption, we conclude the RabE-labeled, organelle-like structures in *P. patens* are likely Golgi, given the observed similarity with previously characterized *A. thaliana* RabE-labeled organelles that co-localize with the Golgi marker sialyltransferase (Zheng et al., 2005; Speth et al., 2009). Moreover, the putative RabE-Golgi structures show similar size to *P. patens’* Golgi and are characteristically excluded from the vesicle-rich apical zone (Furt et al., 2012). In addition, the diffuse cytoplasmic signal and the distinct localization to the growing apical zone are consistent with secretory vesicle localization (Furt et al., 2013; Bibeau et al., 2018). This type of localization is reminiscent of commonly observed localization patterns of F-actin, secretory vesicles, myosin XI/V, and other secretory components across plants and fungi (Bi and Park, 2012; Hepler and Winship, 2015; Riquelme et al., 2018).

Our *in vivo* microscopy and cross-correlation data established that myosin XI and RabE co-localize within the apical spot and are in phase. This data supports our hypothesis that myosin XI and RabE directly interact *in vivo* as observed in Y2H; however, this method and other *in vivo* observation techniques cannot exclude the alternative possibility that myosin XI and RabE are indirectly coupled by another component. To address this limitation, we generated myosin XI mutants with the objective to specifically eliminate binding to RabE and thereby abolish polarized growth. Homology modeling of the hypothesized myosin XI:RabE interaction yielded a putative interface that appears conserved across the large evolutionary distance between plants, yeast, and human. The modeled myosin XI:RabE interface identified structurally conserved residues on the myosin XI tail that were previously demonstrated to be required for polarized growth and Rab binding in yeast (Schott et al., 1999; Pashkova et al., 2006; Jin et al., 2011). Specifically mutating these predicted interface residues abrogated the myosin XI:RabE interaction and eliminated polarized growth. This suggests to us the presence of a similar interface that mediates polarized transport within the last common eukaryotic ancestor of plants, fungi, and animals.

Multiple lines of research support the idea that plant cell division internally recapitulates much of the canonical polarized exocytosis machinery (Fendrych et al., 2010; Mayers et al., 2017). In agreement with previous data independently examining myosin XI and RabE (Zheng et al., 2005; Chow et al., 2008; Yokota et al., 2009; Abu-Abied et al., 2018; Sun et al., 2018), we observe their accumulation at these sites during cell division. However, when we simultaneously observed myosin XI and RabE, we discovered a temporal separation of recruitment, with myosin XI preceding RabE. Intriguingly, the timescale of this separation is similar to myosin XI preceding F-actin and secretory vesicles at the cortical division zone, with myosin XI localizing to the mitotic spindle independent of F-actin (Sun et al., 2018). Microtubule-based transport of myosin XI, similar to myosin VIII (Wu and Bezanilla, 2014), via its cargo-binding domain to the spindle apparatus could explain the uncoupling with RabE and F-actin independence, but this remains to be tested experimentally. Additionally, given myosin XI’s apparent kinetochore localization during chromosome separation, it seems reasonable the cell negatively regulates myosin XI’s “traditional” cargo-binding function to partition polarized transport from chromosome separation to maintain fidelity. We speculate that cells regulate the myosin XI:RabE interaction in a cell-cycle dependent way, possibly through a pathway analogous to yeast, in which RabE localization may be post-translationally controlled via phosphorylation (Lepore et al., 2016). Our observation of RabE rapid colocalization with myosin XI within our observation window at the emerging cell plate may be a consequence of dephosphorylation of RabE. However, this mechanism remains to be determined and is possibly associated with F-actin polymerization at the phragmoplast (Wu and Bezanilla, 2014). Cell-cycle control of polarized secretion, as well as possible interactions with the other plant-specific myosin family, myosin VIII, that is involved in cytokinesis (Wu and Bezanilla, 2014), present fascinating avenues for future research.

Until now, direct evidence for endomembrane-localized myosin XI cargoes was limited to Rab GTPases of the RabC and RabD clades (Hashimoto et al., 2008), and the enigmatic MyoB family (Peremyslov et al., 2013; Stephan et al., 2014). MyoB is a myosin XI effector that localizes to an unknown endomembrane compartment that polarizes to the tip of growing root hairs, as expected for a protein family associated with polarized transport (Peremyslov et al., 2013; Kurth et al., 2017). Unexpectedly, Sadot and co-authors failed to observe MyoB at the growing cell plate (Abu-Abied et al., 2018), another prominent site of polarized transport (McMichael and Bednarek, 2013). Taken together with observations of myosin XI and secretory vesicles at the division site (Sun et al., 2018), this suggests MyoB does not function as myosin’s secretory vesicle receptor. Therefore, we hypothesized a heretofore unidentified myosin XI interactor(s) mediates transport of secretory vesicles. Our work identifying the myosin XI:RabE, establishing the requirement of this interaction for polarized growth, and the observed colocalization at both the growing apex and cortical division zone supports our inference that RabE functions as myosin XI’s secretory cargo receptor.

We suspect the observed myosin XI-2 interaction with RabC and RabD (Hashimoto et al., 2008) is a consequence of the diversification of myosin XIs in angiosperms and does not represent the fundamental and conserved function of myosin XI/V:RabE/Sec4 interaction required for secretory vesicle transport. However, the abundance of myosin XIs in studied angiosperms confounds possible interpretations. We submit that the use of *P. patens* is best suited to parse out the mechanistic functions of myosin XI. Based on our presented observations, we hypothesize *P. patens’* myosin XI maintained the more ancestral functions commonly observed in myosin Vs. Importantly, we believe our findings are translatable to the most similar myosin XIs in angiosperms, myosin XI-K and XI-C/E, which are commonly assumed to be the disproportionately more important myosin XIs for tip growth in root hairs and pollen tubes, respectively. Specifically, we suspect the cargo-binding domain of *Arabidopsis* myosin XI-K associates with RabE. This hypothesis complements the recent discovery in *Arabidopsis* of a putative RabE GEF, the SCD1/SCD2 complex, that localizes to sites of polarized secretion and biochemically interacts with the exocyst (Mayers et al., 2017). We propose a polarized transport model similar to yeast whereby myosin XI transports secretory vesicles by direct interaction with a vesicle-localized RabE GTPase that is activated by its cognate GEF, such as the SCD1/SCD2 complex. With this hypothetical framework, we can inform our experiments to directly test this model and tease out additional mechanistic and dynamic detail, such as the potential coordination of myosin XI, Rabs, and the exocyst to regulate exocytosis in a way that supports rapidly growing cells.

Despite our best efforts, we were unable to reconstitute direct binding of myosin XI to RabE *in vitro*. We attempted traditional equilibrium binding experiments using a quantitative GST pull down assay (Pollard, 2010), as well as using microscale thermophoresis (Wienken et al., 2010). We speculate the myosin XI:RabE dissociation constant is high to facilitate the dynamic fluctuations we observe at the growing tip, which would affect our ability to detect the interaction. Furthermore, it is possible myosin XI interacting with RabE is not a simple binary interaction. Rather, additional components, such as phosphoinositides, could coordinate with membrane-localized RabE to impart a coincidence detection mechanism for specific myosin XI recruitment (Jean and Kiger, 2012).

Here we demonstrate that the RabE subfamily directly interacts with myosin XI, and this interaction is required for proper polarized growth. When coupled with recent discoveries that demonstrate functional homology to the well-characterized yeast system, such as the putative RabE GEF SCD1/SCD2 and exocyst components, our work supports the idea of a myosin XI/V-driven, trans-kingdom mechanism of polarized exocytosis. We anticipate future work will employ this fundamental framework of polarized exocytosis through leveraging experiments in both traditionally “higher” and “lower” plants to build additional complexity that is invariably present in plants, as well as other eukaryotic systems.

## Materials and Methods

### Yeast two-hybrid assay

A Y2H screen was performed using the services of Hybrigenics (France), and we partnered with them to construct their *Physcomitrella patens* library. We cultured moss protonema on PpNO3 and PpNH4 medium under standard conditions (Vidali et al., 2007) for 1-week, then harvested the juvenile tissue and isolated total RNA according to the manufacture’s recommendations (Zymo Research Corp. Cat#R1056). The RNA was immediately frozen, then shipped to Hybrigenics where they constructed the cDNA library. To maximize the probability of discovering true positive interactions, we designed the myosin XI bait fragment according to what we empirically determined to be a soluble protein that encompasses the entire globular tail domain and a piece of the coiled-coil. This MyoXIa-CCT bait fragment corresponds to residues 1061-1536, based on full-length myosin XI protein sequence. MyoXIa-CCT bait fragment was cloned into pB27 (Hybrigenics) and was screened by 132 million pairwise interactions. Positively interacting pairs were identified by growth on SD -Trp -Leu -His. From this deep screen, we recovered the full-length RabE14 coding sequence.

We validated this interaction by repeating the pairing of MyoXIa-CCT and full-length RabE14. We obtained the bait and prey plasmids from Hybrigenics and transformed them into their corresponding Y2H strains, L40 and Y187, respectively. Additional RabEs were inserted into the Y2H prey vector via restriction-based cloning using BamHI and NotI. The MyoXIa-CCT bait was mutagenized with the Q5 Site-Directed Mutagenesis Kit (New England BioLabs) to create the V1418R and V1422R mutants. All constructs were sequence verified and all necessary primers are in supplemental table 1. Bait and prey pairs were mated, then grown in SD -Trp -Leu until ∼2OD. All strains were normalized to 0.3OD, then serially diluted and spotted on SD -Trp -Leu, and SD -Trp -Leu-His plates.

### Construction of fluorescently tagged RabE moss lines

Creation of the 3xmEGFP tagged RabE12 construct occurred by a two-element LR Gateway reaction (ThermoFisher) of entry clones pENT-L1-3xmEGFP-L5r and pENT-L5-RabE12-L2, and the destination vector pTHUbi-gate (Vidali et al., 2007). The RabE12 entry clone was generated by amplification of cDNA using primers AttB2-PpRabE12Rev and AttB5-PpRabE12For (Table S1) and a BP reaction with the resulting PCR product and pDONR 221 P5P2. cDNA was created from RNA isolated from 1-week old moss protonemal tissue using the SuperScript III Reverse Transcriptase (ThermoFisher) according to manufacturer’s protocol. The 3xmCherry-RabE14 construct was generated in a similar manner using RabE14 cDNA, 3xmCherry entry clone, and the destination vector pTKUbi-gate (Wu and Bezanilla, 2014). The moss expression construct pTHUbi-gate:3xmEGFP-RabE12 was transformed into the Gransden wild-type laboratory strain (Liu and Vidali, 2011) and selected for stable transformants. pTKUbi-gate:3xmCherry-RabE14 was transformed into a previously created line that contains myosin XIa endogenously tagged with 3xmEGFP (Sun et al., 2018), as well as wild-type. This procedure yielded multiple independent transformants that were individually screened for fluorescence.

### Live-cell confocal imaging and kymograph analysis

All moss tissue was cultured as previously described (Furt et al., 2013) to enable the use of high-resolution optics to capture fluorescent dynamics within individual growing caulonemal cells. All moss lines were imaged with a SP5 confocal microscope (Leica) using the 488 nm and 561 laser lines at 5% power, with the emission bandwidth gated to 499-547 nm light for GFP and 574-652 nm for mCherry. For imaging of cell division, all images (512×512) were acquired simultaneously at 400 Hz Hz using a HCX-PL-Apo, 63x, NA1.4 lens (Leica) with a zoom of 6 and a bit depth of 12-bit. All images were single medial slices with the confocal aperture at 1 AU, taken at 10 second intervals, and a line average of 2 was used to increase signal to noise. Images were processed by Gaussian blurring (Sigma radius 1), background subtracted (200-pixel ball radius), and contrast enhanced (normalized using stack histogram and 0.1% saturated pixels).

For long-term imaging required for kymograph analysis, all images (1024 x 256 pixels) were acquired simultaneously at 400 Hz using with a zoom of 3. Nine optical slices, 1 µm apart, were collected using at five-second intervals to capture the entire volume of the apical cytosol. Total laser power was maintained at 5% for both laser lines. The confocal aperture was opened to 2 AU. For creation of live-cell movies of growing tips, all images were post-processed with bluring (Sigma radius 1) and the red channel was contrast enhanced (normalized per frame using 0.2% saturated pixels) to adjust for acquisition photobleaching.

For cross-correlation analysis, all data were processed and analyzed in an analogous manner to previous work (Furt et al., 2013). To reduce manual ROI selection and to ensure proper ROI orientation for kymograph analysis, as well as to facilitate data processing and visualization, we implemented a custom MATLAB (MathWorks) routine that is available upon request. For creation of live-cell movies of growing tips, all

### Phylogenetic analyses

The full-length clone identified in our Y2H screen was categorized as RabE14 in accordance with its sequence agreement with the predicted cDNA of gene Pp3c6_11710 (*Physcomitrella patens* v3.3 (Lang et al., 2018)). We then used the predicted protein sequence and BLAST to analyze the *P.patens* proteome using Phytozome (http://phytozome.jgi.doe.gov), as well as *C. subellipsoidea*, *C. reinhardtii*, *V. carteri*, and *A. thaliana*, with an expectation threshold of E = 10^-4^ and BLOSUM62 substitution matrix. All hits were filtered to allow only putative RabE protein sequences for tree construction. We determined if a given sequence was a RabE using the “Rabifier” (rabdb.org) classification pipeline (Diekmann et al., 2011; Surkont et al., 2017) that predicts if the query sequence is a Rab, and if so, groups it into the most likely Rab subfamily based upon canonical subfamily sequence motifs (Pereira-Leal and Seabra, 2000). We added well-characterized Rab8 proteins Sec4 and Rab8a to observe eukaryotic divergence, as well as a known non-Rab8, Ypt1, to function as the outgroup. Sequences were aligned using MUSCLE (Edgar, 2004) under default parameters. The phylogenetic tree was constructed under the ML information criterion using PhyML (Guindon et al., 2010), model selection was performed using the Akaike Information Criterion (Akaike, 1974), and 100 bootstraps were executed for statistical branch support. This process was facilitated through use of the ATGC PhyML web server (Guindon et al., 2005).

### Homology modeling and interface prediction

Homology models of the *P.patens* myosin XI cargo-binding domain (MyoXI-CBD), also referred to as the globular tail domain in the literature, and RabE14 were generated using the SWISS-MODEL server (Waterhouse et al., 2018). The MyoXI-CBD model was created using residues 1131-1537 from full-length myosin XIa and supplying the structure of yeast Myo2p (PDB:2f6h) as the template. The RabE14 active and inactive models consist of the entire protein sequence and used the Sec4 GTP-bound (PDB: 1g17) and GDP-bound (PDB: 1g16), respectively.

Subsequent molecular visualization and manipulation was performed using UCSF Chimera (Pettersen et al., 2004). MyoXI-CBD and active RabE14 were structurally aligned with the myosin Vb-Rab11a co-crystal structure (PDB: 4lx0) using the MatchMaker tool with default parameters. The template was removed, and putative interface residues were extracted through identification of atoms whose respective van der Waals surfaces were within 1 Å. This process was repeated with Myo2p and active Sec4 to assess the ability of the method to recapitulate a known myosin V:Rab8-like interaction, irrespective of the template containing a Rab11 member.

We also employed the template-based docking algorithm, PRISM (Tuncbag et al., 2011; Baspinar et al., 2014). With this approach, a predicted interface is generated based on mapping template-to-target interfaces, thereby bypassing other structural elements unrelated to the interface. We used our homology models as the two target proteins and specified the co-crystal structure of human myosin Vb and active Rab11a as the template. Predicted residue interactions were visualized with a contact map (Fig. 2e). Solvent accessible area was determined using UCSF Chimera and a default probe radius of 1.4 Å.

### RNAi growth assay of myosin XI-CBD mutants

Myosin XI-CBD mutant expression constructs were created via site-directed mutagenesis of wild-type myosin XIa-CBD (Table S1). The mutant and wild-type CBDs were concatenated with the myosin XIa head, neck, and coiled-coil domains through an LR reaction (ThermoFisher) into the moss expression construct, pTHUbi-gate. The full-length myosin XI mutant constructs leave a small linker between the two fused pieces, so we created an otherwise wild-type control construct to test for possible deleterious effects of the linker fragment, named ‘*P.p.* MyoXI.’ The myosin XI-CBD expression constructs were co-transformed (Liu and Vidali, 2011) with a previously developed myosin XI 5’UTR silencing construct (Vidali et al., 2010) to evaluate the capacity of the mutant MyoXI-CBDs to rescue polarized growth. Plants were imaged 1-week post-transformation for polarized growth and size defects (Vidali et al., 2007; Bibeau and Vidali, 2014).

### Myosin XI protoplast expression

The mutant and WT myosin XIa expression constructs were reamplified with primers containing attB1 and attB5 sites (Table S1) to facilitate c-terminal tagging with three tandem copies of monomeric enhanced GFP (3mEGFP). Entry clones of myosin XI (WT and mutants) and 3mEGFP were cloned into a moss expression plasmid with LR clonase (Thermo Fisher), yielding pB1-MyoHNC-B5-gtail-*mutation*-B5-3mEGFP-B2. Constructs were transformed into moss protoplast as previously described (Vidali et al., 2007), except transformed protoplasts were left in suspension in liquid plating medium, instead of being cultured on solid medium.

One day post-transformation, protoplasts were removed from the growth chamber and centrifuged at 250 *g* for five minutes. The supernatant was removed, such that 1ml remained, and the protoplasts were gently resuspended. Slides for imaging were prepared by affixing an approximately 22×22mm square of Parafilm, with a circle cut out of the center, to a glass microscope slide using heat. Approximately 65 µl of protoplasts were transferred to the circle bounded by Parafilm on the slide, then sealed with a 22×22mm glass coverslip. Images of fluorescent protoplasts were acquired using the same equipment described by Bibeau and Vidali (Bibeau and Vidali, 2014).

In brief, protoplasts were imaged using three separate filters: chlorophyll channel (480/40 bandpass excitation filter, 505 longpass dichroic, 510 longpass emission); standard GFP filter (for GFP expression); standard dsRed filter (dead cells/background subtraction). All three images were imported into ImageJ, contrast enhanced, then the dsRed signal was subtracted from the GFP channel to avoid quantification of dead protoplasts. The red channel was used to visualize the protoplast morphology and was manually segmented using the magic wand tool and ROI manager. The mean gray values for all ROIs were measured on the GFP channel and exported. All fluorescent values were filtered based on the fluorescence of the “no DNA” condition—the maximum value obtained within the “no DNA” condition was used as a threshold for all other conditions for that respective day to discard untransformed or non-expressing protoplasts. The remaining values were averaged by date of transformation for each condition and tested for statistical differences.

### Myosin XI-CCT purification

The myosin XIa-CCT *E. coli* expression construct was created by primers (Table S1) that amplified residues 1061-1537 and then cloned into the pETDuet vector using restriction enzymes BamHI and HindIII. Myosin XI mutants were generated for purification using the Q5 Mutagenesis Kit (New England BioLabs) using the previously created mutagenesis primers for protoplast expression and the wild-type MyoXIa-CCT *E. coli* expression construct as the template.

Myosin XIa-CCT expression constructs were transformed into BL21 (DE3) *E. coli*, transferred to liquid culture and grown until OD600 ∼0.6, then shifted to 15oC and induced with 0.1 mM IPTG after cooling. Cells were incubated overnight at 15oC. Harvested cells were resuspended (∼20 ml lysis buffer/L of culture) in cold lysis buffer (50 mM NaH2PO4 pH 8.0, 300 mM NaCl, 10mM imidazole, 10% (v/v) glycerol, fresh 5mM β-mercaptoethanol, 1mM PMSF, DNAse, and 1 cOmplete Protease Inhibitor Cocktail tablet (Roche Diagnostics) added prior to lysis, then lysed using a microfluidizer M-110S (Microfluidics) at 80 psi. The cleared lysate was batch bound to Ni-NTA agarose beads (Qiagen) for 1 h at 4oC, then packed via gravity flow. The Ni-NTA column was washed (50 mM NaH2PO4 pH8.0, 300 mM NaCl, 20 mM imidazole, 10% (v/v) glycerol, fresh 5 mM β-mercaptoethanol), then eluted with the same buffer plus 250 mM imidazole. Fractions containing protein were pooled and subjected to an anion exchange purification step on a MonoQ 10/10 column (GE LifeSciences), controlled using an ÄKTA pure FPLC (GE LifeSciences).

### Western blot analysis

Y2H strains were analyzed for MyoXIa-CCT expression by picking individual colonies from SD -trp -leu plates, growing to 1 OD600 unit, and extracting total protein via a NaOH and SDS-PAGE buffer treatment (Kushnirov, 2000). Total protein was run on a 12% SDS-PAGE gel, transferred to a PVDF membrane, cut, then blotted against myosin XI-CCT (1:10,000--∼80kDa) using a custom-made antibody, and yeast ADH1 (1:1,000--∼40kDa) (Abcam:ab34680) as a loading control. The myosin XIa-CCT antibody was developed against a 6xHis fusion of the myosin XIa-CCT (Capralogics, Inc. Hardiwick, MA).

### Statistical Analyses

All statistical analyses were performed using GraphPad Prism. All error bars display the standard error of the mean. For the transient RNAi experiment, statistical significance was determined by a one-way ANOVA with a post-hoc Tukey test (P < 0.01). For the moss protoplast experiment, a one-way ANOVA (P < 0.05) failed to identify a significant difference between any experimental groups.

## Author Contributions

R.G.O., F.F., L.V., and M.M. designed the research. R.G.O., F.F., E.L.W., E.M.A., J.M.G., S.E.C., M.L.D., and A.M.B. conducted experiments and analyzed the data. R.G.O., L.V., and M.M. interpreted the data. R.G.O. wrote the initial draft of the manuscript and M.M and L.V. edited the draft. All authors reviewed and approved the final version of the manuscript.

## Acknowledgements

The authors are grateful to Jeffrey P. Bibeau for developing the XCorr script in Matlab and Miye Jacques for creating the myosin XIa-CCT *E. coli* expression construct. We would also like to thank Sebastian Bednarek for critical feedback on this work. This work was supported by NSF grant MCB-1253444 to L.V. and by NIH grant #1R01 GM068803 to M.M.

