## Supplemental Figures for "Myosin XI Interacting with a RabE GTPase Is Required for Polarized Growth"

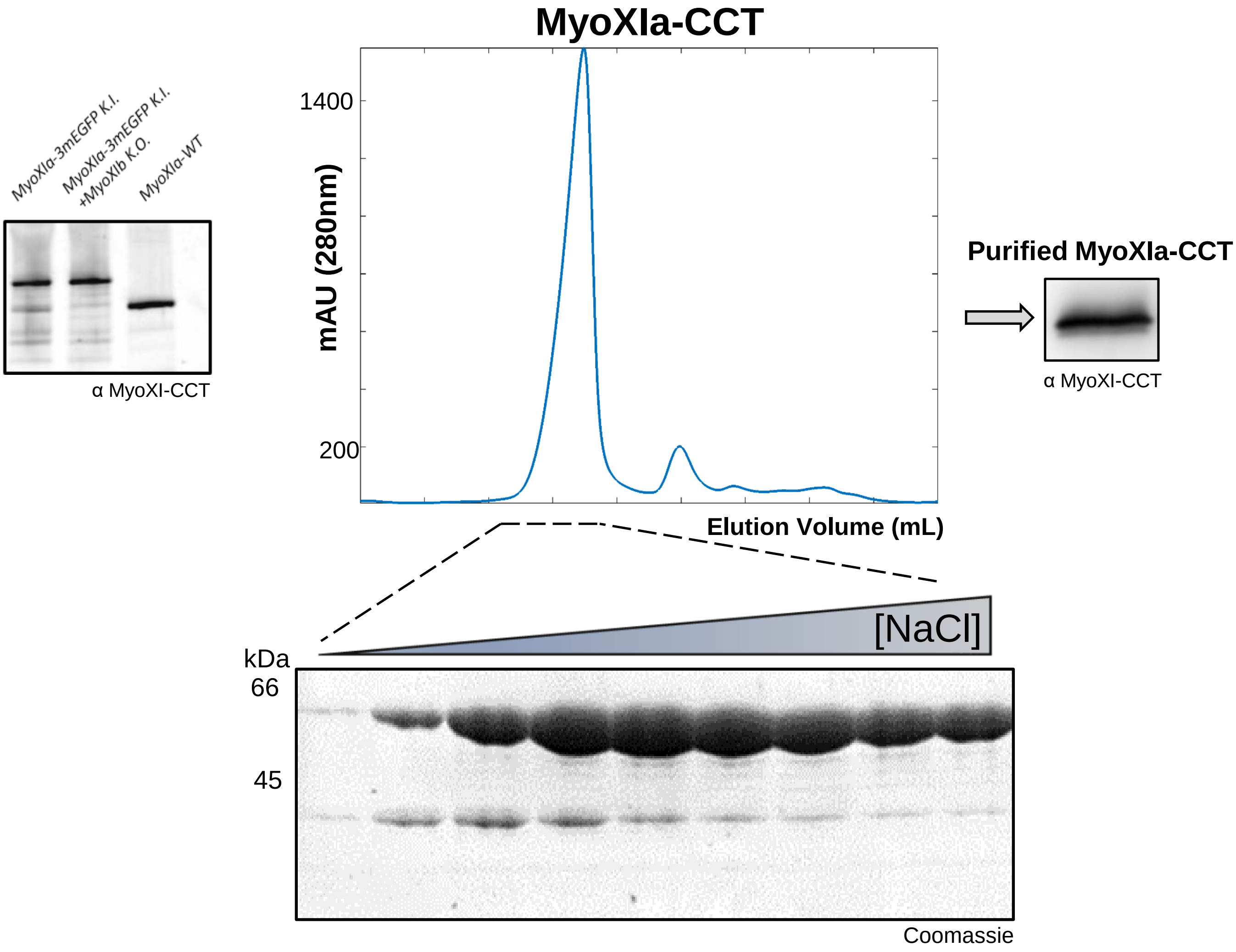


**Supplemental Figure 1. Purification of Myosin XIa-CCT and Validation of Myosin XIa Antibody.**

(Supports Figure 1.)

Our in-house developed antibody to MyoXIa-CCT was tested against lysates of different moss lines, as well as purified MyoXIa-CCT. The western blot shows the expected size shift between endogenously tagged myosin XI fused to 3xmEGFP and wild-type myosin XI from moss protein extracts. The Coomassie-stained SDS-PAGE gel shows purified fractions of MyoXIa-CCT in correspondence to the anion-exchange salt elution profile. The fractions were pooled, buffer exchanged, and tested against our MyoXIa-CCT antibody through western blot.

A

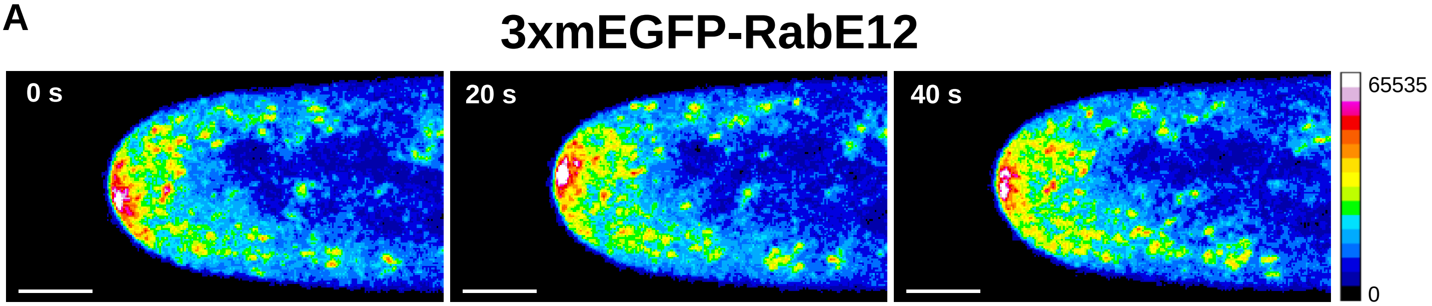


B

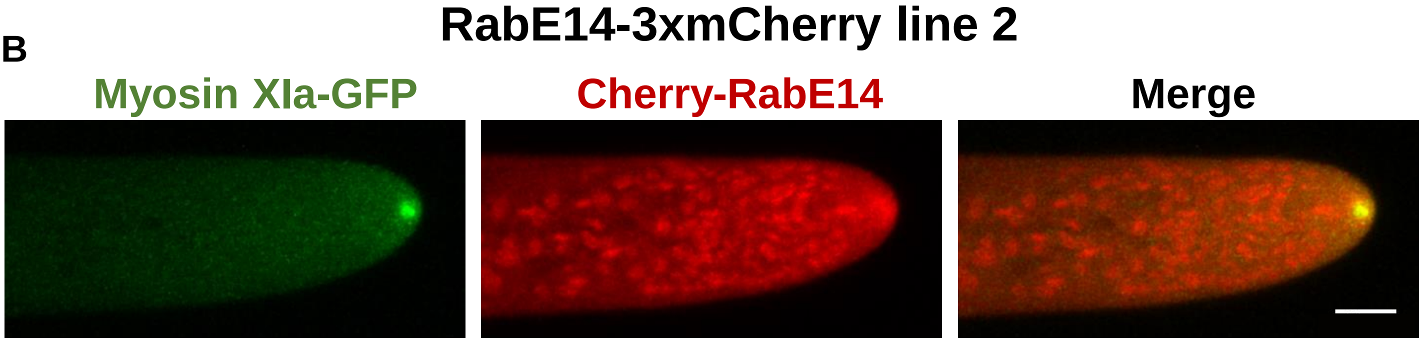


**Supplemental Figure 2. Subcellular Localization of RabE12 and Independent RabE14 Lines**. (Supports Figure 2.)

(A) Subcellular localization of 3xmEGFP-RabE12 by confocal microscopy. Images are maximum projections of 3 confocal slices to visualize the medial volume of the cell and acquired at 5-sec intervals. Selected images from a time series are shown using a 16-color look-up table with the corresponding pixel value range shown on the right.

(B) Subcellular localization of 3xmCherry-RabE14 and MyosinXIa-3xmEGFP from an independently isolated line. Scale bar, 5 µm

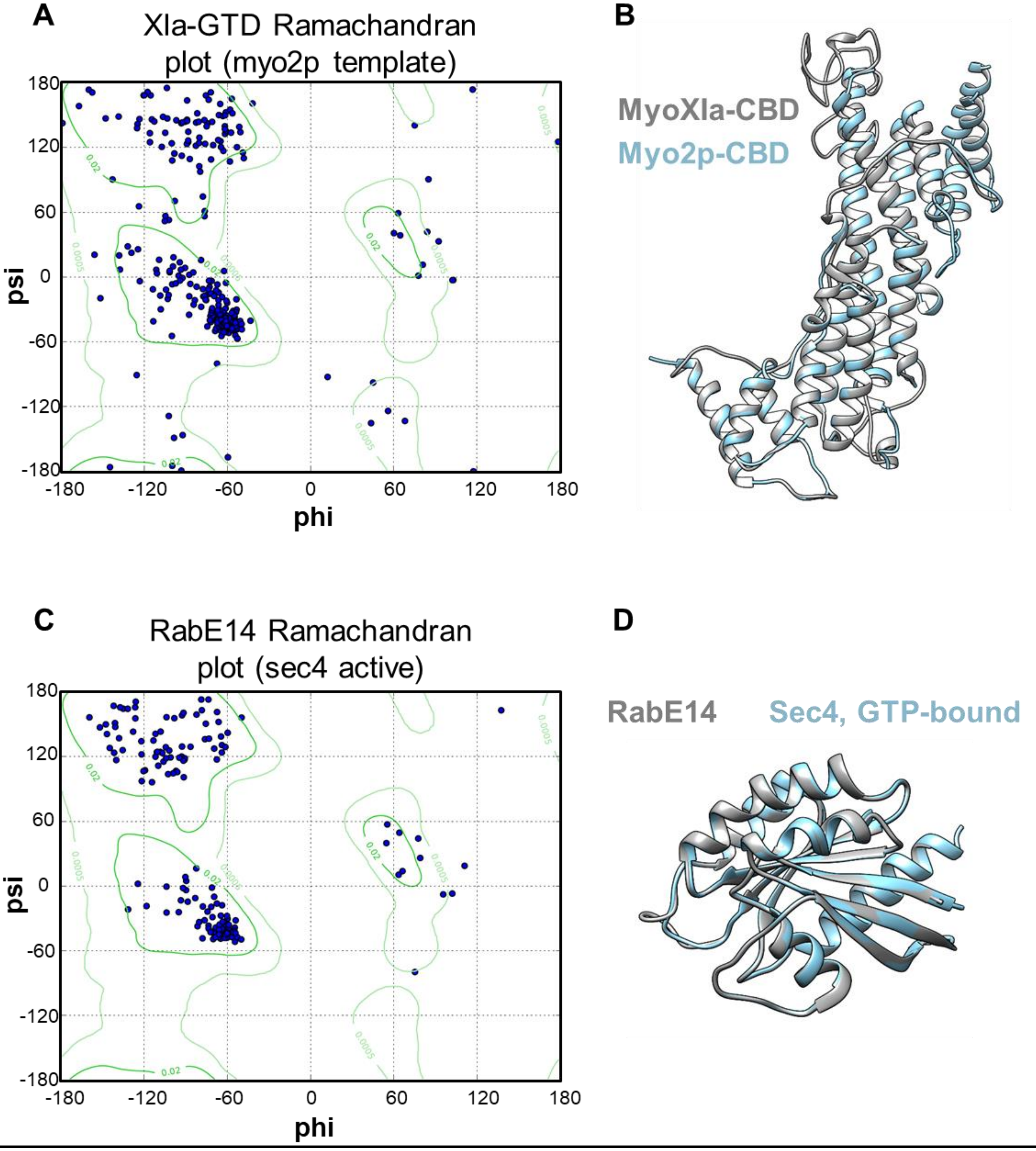


**Supplemental Figure 3. Ramachandran plots and average RMSDs for MyoXIa-CBD and RabE14 homology models**

(Supports Figure 4.)

(A-D) Both MyoXIa-CBD and RabE14 homology models used for interface prediction exceed the conventional cutoff of >95% of all non-glycine residues must occupy the energetically allowed regions within the Ramachandran map to conclude good stereochemical quality.

(A) Eight non-glycine residues of MyoXIa-CBD are located outside of these regions (corresponding to ~98% within).

(B) The average RMSD of MyoXIa-CBD is 0.291 Å.

(C) Four non-glycine residues of active RabE14 located outside (corresponding to ~98% within).

(D) The average RMSD of active RabE14 is 0.134 Å.

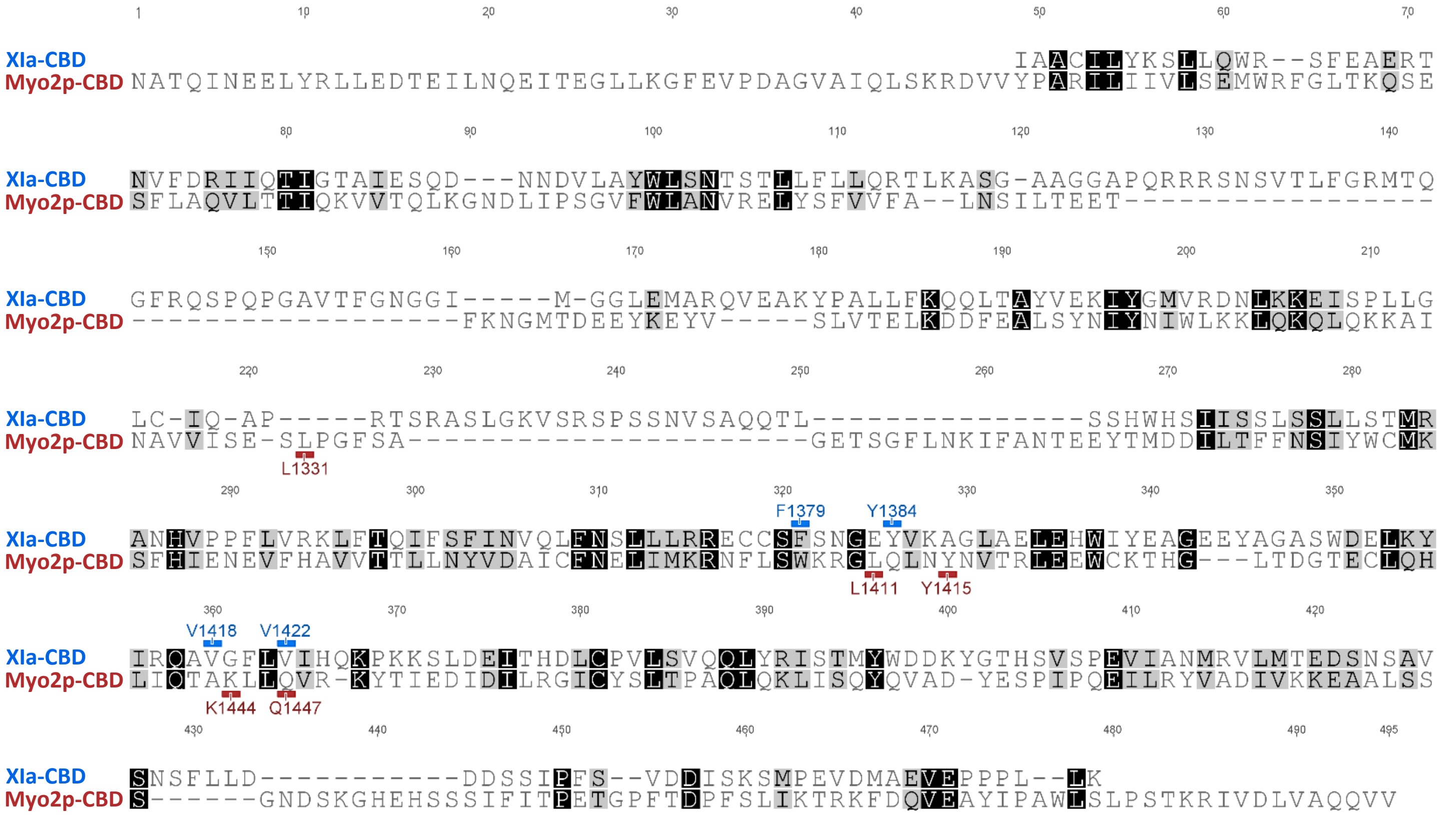


**Supplemental Figure 4. A structure-based sequence alignment of MyoXIa-CBD homology model and Myo2p-CBD crystal structure**

(Supports Figure 5)

Structure-based sequence alignment was performed using UCSF Chimera using our homology model of *P. patens* myosin XI cargo-binding domain and crystal structure of *S. cerevisiae* myosin V cargo-binding domain (Myo2p-CBD), PDB=2f6h.

| Primer Name | Primer Sequence | Entry clone | Use |
|---|---|---|---|
| AttB1 Myo Head Neck For | GGGGACAAGTTTGTACAAAAAAGCAGGCTTAATGGCGACAGCAGGGAATGTA | pL1-MyoHeadNeck-R5 | Phenotype/Complementation |
| AttB5r Myo Head Neck Rev | GGGGACAACTTTTGTATACAAAGTTGTACCCTTGGTCGGTGACAATAC | pL1-MyoHeadNeck-R5 | Phenotype/Complementation |
| AttB5 Myo CC Tail For | GGGGACAACTTTGTATACAAAAGTTGTGTTGTCGAATCGATTCAAAAGCACCG | pL5-MyoCCTail-L2 | Phenotype/Complementation |
| AttB2 Myo CC Tail Rev | GGGGACCACTTTGTACAAGAAAGCTGGGTACTAAGAATCTGGTTGTGG | pL5-MyoCCTail-L2 | Phenotype/Complementation |
| AttB1 Myo Head Neck CC For | GGGGACAAGTTTGTACAAAAAAGCAGGCTTAATGGCGACAGCAGGGAATGTA | pL1-MyoHeadNeckCC-R5 | Phenotype/Complementation |
| AttB5r Myo Head Neck CC Rev | GGGGACAACTTTTGTATACAAAGTTGTCAAAGAGTCTTGATTCTCCTGCT | pL1-MyoHeadNeckCC-R5 | Phenotype/Complementation |
| AttB5 Myo Tail For | GGGGACAACTTTGTATACAAAAGTTGTGCTGCAATGTGTCATGCAAGAT | pL5-MyoTail-L2 | Phenotype/Complementation |
| AttB2 Myo Tail Rev | GGGGACCACTTTGTACAAGAAAGCTGGGTACTAAGAATCTGGTTGTGG | pL5-MyoTail-L2 | Phenotype/Complementation |
| Myo F1379R For | CGTGAGTGTTGCTCACGTAGCAACGGAGAGTAT | pL5-MyoTailF1379R-L2 | Phenotype/Complementation |
| Myo F1379R Rev | TCTCAGCAGCAAACTGTTGAACAGCTGAACATT | pL5-MyoTailF1379R-L2 | Phenotype/Complementation |
| Myo V1418R For | TATATCCGACAAGCACGTGGATTTTTGGTCATTCATC | pL5-MyoTailV1418R-L2 | Phenotype/Complementation/Purification |
| Myo V1418R Rev | CTTGAGCTCATCCCATGACGCTCCAGCATACTC | pL5-MyoTailV1418R-L2 | Phenotype/Complementation/Purification |
| Myo V1422R For | GCAGTTGGATTTTTGCGCATTCATCAAAAGCCA | pL5-MyoTailV1422R-L2 | Phenotype/Complementation/Purification |
| Myo V1422R Rev | TTGTCGGATATACTTGAGCTCATCCCATGACGCTCC | pL5-MyoTailV1422R-L2 | Phenotype/Complementation/Purification |
| Myo W1408R For | TATGCTGGAGCGTCACGGGATGAGCTCAAGTAT | pL5-MyoTailW1408R-L2 | Phenotype/Complementation |
| Myo W1408R Rev | CTCCTCCCCAGCTTCATAAATCCAGTGCTCTAGTTCTG | pL5-MyoTailW1408R-L2 | Phenotype/Complementation |
| Myo Y1384R For | TTTAGCAACGGAGAGCGTGTGAAAGCTGGACTT | pL5-MyoTailY1384R-L2 | Phenotype/Complementation |
| Myo Y1384R Rev | TGAGCAACACTCACGTCTCAGCAGCAAACTGTT | pL5-MyoTailY1384R-L2 | Phenotype/Complementation |
| MyoXIa-CCT WT For | CGCGGATCCGTTGTCGAATCGATTCAAAAGC | n/a | BamHI site for cloning into pETDuet/Purification |
| MyoXIa-CCT WT Rev | CCCAAGCTTCTAAGAATCTGGTTGTGGCATTAG | n/a | HindIII site for cloning into pETDuet/Purification |
| AttB2-PpRabE12 Rev | GGGGACCACTTTGTACAAGAAAGCTGGGTACTACGAGCAGCAGGAACTAGA | pL5-RabE12-pL2 | Subcellular localization of RabE12/3xmEGFP tagging |
| AttB5-PpRabE12 For | GGGGACAACTTTGTATACAAAAGTTGTGATGGCCGCAGGTGGATCAAGA | pL5-RabE12-pL2 | Subcellular localization of RabE12/3xmEGFP tagging |
| AttB5-PpRabE14 For | GGGGACAACTTTGTATACAAAAGTTGTGATGGCGACAAGAGCCC | pL5-RabE14-pL2 | Subcellular localization of RabE14/3xmEGFP tagging |
| AttB5-PpRabE14 Rev | GGGGACCACTTTGTACAAGAAAGCTGGGTACTATGAGCAGCAAGAGCCAGA | pL5-RabE14-pL2 | Subcellular localization of RabE14/3xmEGFP tagging |
| RabE11CDS_NotI_Forward | TAAGCAGCGGCCGCAATGGCCGCAGGTGGATC | y2hPrey-RabE11 | RabE11 full-length in Y2H prey vector |
| RabE11CDS_BamHI_Reverse | TGCTTAGGATCCTCAAGAGCAACAGGAGCTACC | y2hPrey-RabE11 | RabE11 full-length in Y2H prey vector |
| RabE15CDS_NotI_Forward | TAAGCAGCGGCCGCAATGGCGACAAGAGCTCGG | y2hPrey-RabE15 | RabE15 full-length in Y2H prey vector |
| RabE15CDS_BamHI_Reverse | TGCTTAGGATCCCTATGAGCAGCAAGAGCCAGA | y2hPrey-RabE15 | RabE15 full-length in Y2H prey vector |
| Y2Hprey_Forward | AATACCACTACAATGGAT | n/a | Sequencing primer for Y2H prey vector |
| Y2Hprey_Reverse | TCTAGACACTAGCTACTC | n/a | Sequencing primer for Y2H prey vector |

**Supplemental Table 1.** Primers used in this study.

| Name | Description | Sequence Length |
|---|---|---|
| Volvox carteri | Vocar.0012s0034.1 (Phytozome) | 217 |
| S.c. Ypt1 | P01123 (Uniprot) | 206 |
| Coccomyxa subellipsoidea | 26393 (Phytozome) | 210 |
| Chlamydomonas reinhardtii | Cre15.g641800.t1.2 (Phytozome) | 183 |
| H.s. Rab8a | P61006 (Uniprot) | 207 |
| A.t. RabE1E | At3g09900 (Phytozome-TAIR10) | 218 |
| A.t. RabE1D | At5g03520 (Phytozome-TAIR10) | 216 |
| A.t. RabE1B | AT5G59840 (Phytozome-TAIR10) | 216 |
| A.t. RabE1A | At3g53610 (Phytozome-TAIR10) | 216 |
| A.t. RabE1C | At3g46060 (Phytozome-TAIR10) | 216 |
| P.p. RabE15 | Pp3c5_21410 (Phytozome-Pp v3.3) | 215 |
| P.p. RabE13 | Pp3c25_7430 (Phytozome-Pp v3.3) | 216 |
| P.p. RabE14 | Pp3c6_11710 (Phytozome-Pp v3.3) | 215 |
| P.p. RabE11 | Pp3c16_17460 (Phytozome-Pp v3.3) | 216 |
| P.p. RabE12 | Pp3c16_17470 (Phytozome-Pp v3.3) | 216 |
| S.c. Sec4 | P07560 (Uniprot) | 215 |

**Supplemental Table 1. Gene IDs of sequences used in phylogenetic tree construction.**
(Supports Figure 4.)
