## Supplemental Movie Legends for "Myosin XI Interacting with a RabE GTPase Is Required for Polarized Growth"

**Movie 1** Confocal live-cell imaging of the 3xmCherry-RabE14 and MyosinXIa-3xmEGFP transgenic line used for analysis in Figure 2. Images were acquired as z-stacks of 9 confocal slices, at a rate of 5 seconds per frame. The confocal slices were maximum projected. The movie frame rate is 20 frames per second. Time is indicated as minutes:seconds, and the scale bar is 5 µm.

**Movie 2** Confocal live-cell imaging of the 3xmEGFP-RabE12 transgenic shown in Supplemental Figure 2a. Images were acquired as z-stacks of 6 confocal slices, at a rate of 5 seconds per frame. The medial 4 confocal slices were maximum projected. The movie frame rate is 20 frames per second. Time is indicated as minutes:seconds, and the scale bar is 5 µm.

**Movie 3** Confocal live-cell imaging of the 3xmCherry-RabE14 and MyosinXIa-3xmEGFP transgenic line shown in Figure 3. Images are single medical confocal sections acquired at a rate of 10 seconds per frame. The first frame occurs approximately 1-minute post nuclear envelope breakdown. The movie frame rate is 15 frames per second. Time is indicated as minutes:seconds, and the scale bar is 5 µm.
